# The sulfur cycle connects microbiomes and biogeochemistry in deep-sea hydrothermal plumes

**DOI:** 10.1101/2022.06.02.494589

**Authors:** Zhichao Zhou, Patricia Q. Tran, Alyssa M. Adams, Kristopher Kieft, John A. Breier, Rupesh K. Sinha, Kottekkatu P. Krishnan, P. John Kurian, Caroline S. Fortunato, Cody S. Sheik, Julie A. Huber, Meng Li, Gregory J. Dick, Karthik Anantharaman

## Abstract

In globally distributed deep-sea hydrothermal vent plumes, microbiomes are shaped by the redox energy landscapes created by reduced hydrothermal vent fluids mixing with oxidized seawater. Plumes can disperse over thousands of kilometers and are complex. Their characteristics are determined by geochemical sources from hydrothermal vents, e.g., hydrothermal inputs, nutrients, and trace metals. However, the impacts of plume biogeochemistry on the oceans are poorly constrained due to a lack of integrated understanding of microbiomes, population genetics, and geochemistry. Here, we use microbial genomes to understand links between biogeography, evolution, and metabolic connectivity, and elucidate their impacts on biogeochemical cycling in the deep sea. Using data from 37 diverse plumes from 8 ocean basins, we show that sulfur metabolism defines the core microbiome of plumes and drives metabolic connectivity. Amongst all microbial metabolisms, sulfur transformations had the highest MW-score, a measure of metabolic connectivity in microbial communities. Our findings provide the ecological and evolutionary basis of change in sulfur-driven microbial communities and their population genetics in adaptation to changing geochemical gradients in the oceans.

## Main

Hydrothermal vents are abundant and widely distributed across the deep oceans. The mixing of hot hydrothermally-derived fluids rich in reduced elements, compounds, and gasses, with cold seawater forms hydrothermal plumes^1, 2^. Usually, plumes rise up to hundreds of meters from the seafloor and can disperse over hundreds to thousands of kilometers through the pelagic oceans^3^. Surrounding microbes migrate into the plume and thrive on substantial reductants as the energy sources, making plumes ‘hotspots’ of microbial activity and geochemical transformations^1, 2^. Plumes constitute a relatively closed ecosystem that depends on chemical energy-based primary production and is mostly removed from receiving inputs of energy from the outside^4, 5^. Thus, plumes serve as an ideal natural bioreactor to study the processes and links between microbiome and biogeochemistry and the underlying ecological and evolutionary basis of microbial adaptation to contrasting conditions between energy-rich plumes and the energy-starved deep-sea^2^.

The most abundant energy substrates for microorganisms in hydrothermal plumes include reduced sulfur compounds, hydrogen, ammonia, methane, and iron^2^. Amongst these, sulfur is a major energy substrate for diverse microorganisms in plumes across the globe^2, 6, 7, 8^. Sulfur transformations in plumes are dominated by oxidation of reduced sulfur species, primarily hydrogen sulfide and elemental sulfur. The metabolic pathways include oxidation of sulfide to elemental sulfur (*fcc, sqr*), oxidation of sulfur to sulfite (*dsr, sor*, and *sdo*), disproportionation of thiosulfate (*phs*) to hydrogen sulfide and sulfite, disproportionation of thiosulfate to elemental sulfur and sulfate (*sox*), thiosulfate oxidation to sulfate (*sox, tst*, and *glpE*), and sulfite oxidation to sulfate (*sat, apr*)^7, 9, 10, 11^. Complete oxidation of sulfur would involve oxidation of hydrogen sulfide all the way to sulfate. However, recent observations in other ecosystems indicate that individual microbes rarely possess a full set of the complete sulfide/sulfur oxidation pathway^10, 12^, instead individual steps are distributed across different community members. This likely suggests that sulfur oxidation is a microbial community-driven process that is dependent on metabolic interactions, and asks for revisiting sulfur metabolism and biogeochemistry based on a holistic perspective of the entire community.

Recent microbiome-based ecological studies have focused on elucidating a genome-centric view of ecology and biogeochemistry^7, 10, 12, 13, 14, 15^. This approach has expanded our understanding of microbial diversity associated with specific energy metabolisms, including sulfur transformations in hydrothermal plumes, the deep sea, and beyond^7, 14, 16, 17, 18, 19^. However, the dynamics and microdiversity of the plume microbiome, and relevant biogeochemical impacts remain relatively underexplored^20, 21, 22, 23, 24^. Understanding how environmental constraints and selection shape the microdiversity and the genetic structure of plume microbial populations after migration from background seawater can provide fundamental insights into adaptation mechanisms. These insights can also inform future predictions of microbial responses to the changing oceans.

Here, we characterized the ecological and evolutionary bases of the assembly of the plume microbiome, and their strategies for sulfur cycling-based energy metabolisms. First, we studied globally distributed hydrothermal plume datasets to define a core plume microbiome. We followed this up with synthesis of genome-resolved metagenomics, metatranscriptomics, and geochemistry from three hydrothermal vent sites (Guaymas Basin, Mid-Cayman Rise, and Lau Basin) to unravel community structure and functional links to biogeochemistry, metabolic connectivity within plume and deep-sea communities, and microdiversity in abundant microbial populations. We demonstrate that plume microbiomes have a distinctive community composition and function, that is adapted towards energy conservation, metabolic interactions, and stress response.

## Results

We used publicly available microbiome data from hydrothermal vent plumes across the globe to (1) define the core plume microbiome, (2) investigate plume microbiome structure, function, and activity, and (3) identify links between plume microbiomes and geochemistry. To investigate the core microbiome, we studied publicly available 16S rRNA gene datasets of hydrothermal plumes (*n* = 37) and background deep-sea samples (*n* = 14) from eight ocean basins across the globe. To study the microbiome structure, function, and activity, we reconstructed metagenome-assembled genomes (MAGs) from three hydrothermal vent sites (containing both plume and background samples from Guaymas Basin, Mid-Cayman Rise, and Lau Basin). We also mapped paired metatranscriptomes from the same sites for some samples (Fig. 1, Fig. S1, and Supplementary Data 1). To study links between biogeochemistry and the microbiome, we analyzed paired geochemical data from the above three hydrothermal vent sites. To provide clarity on the plume and background samples, and DNA/cDNA libraries used in this study, we provided a schematic diagram describing the locations of all samples in the context of a hydrothermal vent system (Fig. S1).

**Fig. 1.**
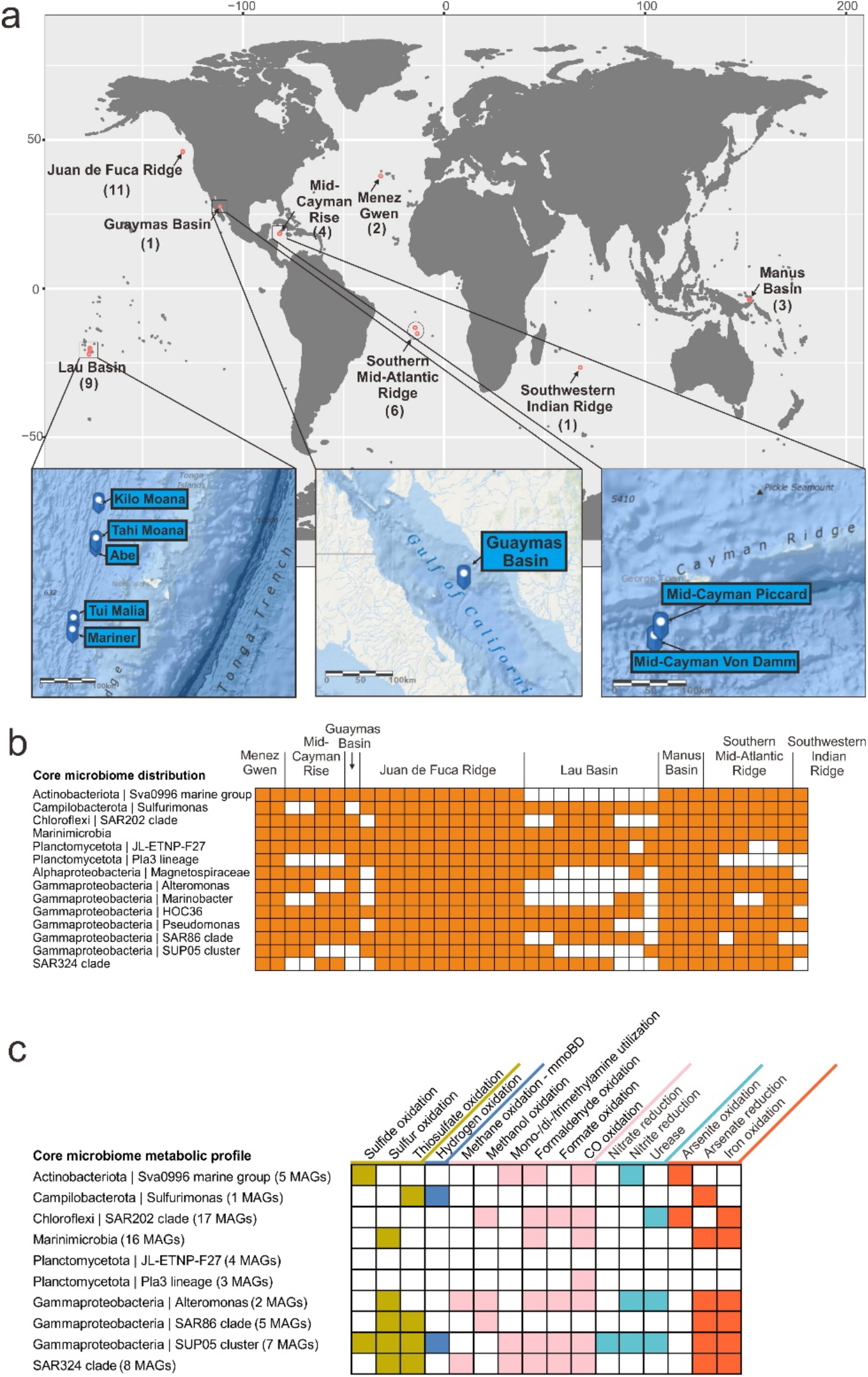
Sampling sites, distribution, and metabolic profile of the core plume microbiome. **a** Sampling site maps of hydrothermal plume samples from which the 16S rRNA gene datasets were sourced. Numbers in brackets indicate dataset quantities. Three hydrothermal sites that have metagenome and metatranscriptome datasets in this study were specifically represented by inset maps. Ocean maps were remodified from ArcGIS online map (containing layers of “World Ocean Base” and “World Ocean Reference”; https://www.arcgis.com/). **b** Membership and distribution of the core plume microbiome. Heatmap shows the presence/absence of core plume microbial groups (tracing back to known taxonomic ranks from the genus-level taxa) in 37 hydrothermal plume 16S rRNA gene datasets across the world. **c** Metabolic profile of the core plume microbiome. From this study, MAGs that have 16S rRNA genes affiliated to the core plume microbiome were used as representatives (numbers labeled in brackets). This subpanel shows the presence or absence of metabolic potential associated with sulfur, carbon, nitrogen, hydrogen, and metal biogeochemical transformations.

### Defining the core hydrothermal plume microbiome

To identify and study the core hydrothermal plume microbiome, we used 16S rRNA gene datasets from 51 hydrothermal plume and background deep-sea samples spread across eight ocean basins (Supplementary Data 2). Biogeographic patterns were delineated by Unifrac metrics of distance and PCoA-based ordination. Sample location had a stronger influence on biogeographic patterns than sample characteristics (plume/background) (Fig. S2, S3). Unweighted Unifrac PCoA plots indicated that paired plume/background deep-sea samples within the same site were closely correlated (Fig. S3). As revealed previously^2, 25, 26^, this supports the understanding that the hydrothermal plume environment has its main constitutional microorganisms derived from surrounding seawaters, with dispersal limitation having little effects locally.

We then identified genus-level taxa significantly distributed in plumes with high prevalence and relative abundance. The core plume microbiome consists of 14 microbial groups (Fig. 1a, b) as revealed from the 37 plume datasets with a cutoff of being distributed in at least two third of all plume datasets and having at least 1% relative abundance on average. By choosing MAGs reconstructed from this study that were affiliated to the same taxa, we characterized metabolic profiles for the core plume microbiome which demonstrated highly versatile metabolic potential for utilizing various plume substrates^2^, including HS^-^, S^0^, H_2_, CH_4_, methyl-/C_1_ carbohydrates, arsenite, and iron (Fig. 1c). Most plume microorganisms are of seawater origin, consistent with prior reports^26^ (Supplementary Table 1). We also observed a small number of seafloor/subsurface dwelling and endosymbiotic microorganisms that might be entrained in plumes^2, 27^ (Supplementary Table 1). Collectively, our data suggest that sulfur and other reduced organic/inorganic compounds significantly shape the global core plume microbiome that are originally derived from the surrounding seawater.

### Distinctive plume geochemistries influence energy landscapes and promote microbial growth

Previous thermodynamic modeling analyses have reflected energy landscapes for various hydrothermal ecosystems^4, 7, 10, 16^ by representing free energy yields for reactions of various energy sources for microbial metabolism in hydrothermal fluids. Some of them have demonstrated the consistency of thermodynamic modeling and omics-based biogeochemical estimation in individual ecosytems^7, 10, 16^. Here based on geochemical parameters and predicted functions from reconstructed MAGs (Fig. S4, S5, and Supplementary Data 3), we conducted an across-site comparison of thermodynamic modeling and omics-based biogeochemical estimations to reflect the influences of distinctive plume geochemical characteristics on plume microbes. We also conducted growth rate analyses to identify whether microbial energy contributors are promoted with higher growth rates in responding to differing geochemical conditions across plumes. To address these, we first reconstructed plume energy landscapes through thermodynamic modeling (Fig. 2a).

**Fig. 2.**
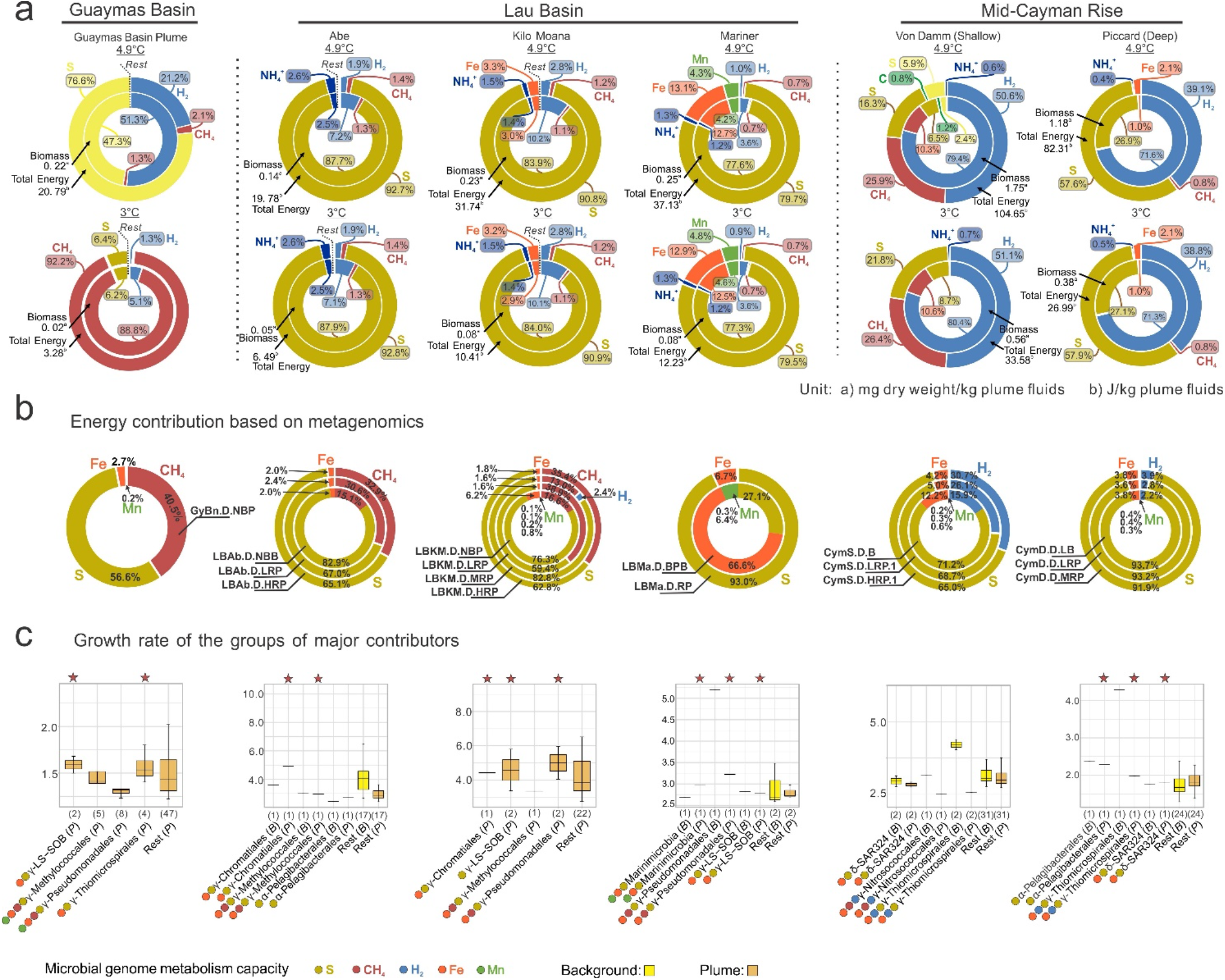
Thermodynamic estimation of available free energies and biomass yields from electron donors, metagenomics-based contribution of electron donors to energy, and growth rates of major microbial contributors. **a** Thermodynamic estimation diagram of available free energy and biomass. For each hydrothermal environment, the contribution fraction of each electron donor species was labeled accordingly in the rings. The total available free energies and biomass were labeled accordingly to individual plumes. Two temperatures (3°C and 4.9°C) were picked to represent *in situ* temperatures in the upper and lower plume. Light yellow represents aerobic sulfur oxidation, dark yellow represents anaerobic sulfur oxidation. **b** Metagenomics-based estimation of energy contribution. Energy contribution for each electron donor was calculated based on metagenomic abundance of each reaction of electron donors and free energy yield of each reaction. The contribution ratio of electron donor species was calculated for individual environments respectively. **c** Growth rate of major microbial contributors in each hydrothermal environment. The y-axis for each barplot indicates the replication rate. The microbial groups starting with “α-”, “γ-”, and “δ-” represent Alphaproteobacteria, Gammaproteobacteria, and Deltaproteobacteria, respectively. Plume microbial groups were colored by dark yellow, background microbial groups were colored by light yellow and they were also all labeled with “(*P*)” or “(*B*)”, respectively. Numbers in brackets indicate MAG numbers in each microbial group. Star-labeled plume microbial groups had higher growth rates than the ‘Rest’ plume microbial groups.

Distinctive geochemical characteristics support the predicted energy landscapes when compared among sites. Methane was the highest in end-member fluids from Guaymas Basin (63.4 mmol/kg)^7^, which supported the dominance of methane oxidation in the Guaymas Basin plume in the thermodynamic model (Fig. 2a), and significant contributions of methane oxidation in metagenomics datasets were also found (∼40.5%) (Fig. 2b). Meanwhile, Lau Basin hydrothermal fluids had high Mn and Fe concentrations (Mn: 3.9-6.3 mmol/kg, Fe: 3.8-13.1 mmol/kg)^28, 29^ in the Mariner hydrothermal field compared to other samples. This manifested in Fe and Mn oxidation contributing the highest fractions (Mn: ∼4-5%, Fe: 13%) in thermodynamic modeling (Fig. 2a) and the highest fractions (Mn: 0.3-6.4%, Fe: 6.7-66.6%) in omics-based estimations of Mariner among all sites (Fig. 2b). Similarly in Mid-Cayman Rise, high hydrogen concentrations in the vent fluids were associated with high contribution of hydrogen oxidation in the model, and in omics-based estimations (Fig. 2a, 2b, Supplementary Table 2). Overall, reduced sulfur is the major energy source as reflected in both thermodynamic modeling and omics-based biogeochemical estimations in all three sites. However, individual plume geochemical conditions vary with diverse minor energy sources, such as iron, methane, and hydrogen, leading to different energy landscapes which are mediated by microbes.

To study whether abundant organisms conducting biogeochemical transformations in each site were also growing actively, we predicted microbial growth rates from metagenomic data using iRep^30^. iRep can use a combination of cumulative GC skews and abundance of metagenomic reads to calculate the difference in read abundance at the origin and terminus of a genome which is a proxy for the replication or growth rate of organism^30, 31, 32^. The results suggest potential associations between growth rates and geochemically-influenced energy landscapes for individual sites (Fig. 2c). A consistent pattern of the abundant microorganisms in plumes having a higher predicted growth rate was also observed in certain sites. For instance, LS-SOB and Thiomicrospirales both had the capacities for sulfur and iron oxidation, and were predicted to have a higher growth rate than other microorganisms in Guaymas Basin plume (Fig. 2c). Similarly, Methylococcales and Chromatiales were the major contributors to iron, methane, and sulfur oxidation in Lau Basin (Abe plume) and their growth rates were higher than other organisms (Fig. 2c). Collectively, we found a consistent pattern demonstrating that the abundant microorganisms also have higher predicted growth rates potentially due to their ability to respond to varying geochemistry in hydrothermal plumes.

### Across- and within-site comparisons for plumes show consistent links between geochemistry, function, and taxonomy

MAGs reconstructed from Guaymas Basin, Mid-Cayman Rise, and Lau Basin hydrothermal vents and corresponding omics-based profiling enabled taxonomic and functional comparisons among the three sites (Fig. S4, S5, and Supplementary Data 3). Across-site analyses of functional traits in MAGs indicate that different functions were significantly enriched in different plumes, e.g., arsenate reduction and long-chain alkane (C_6_ +) degradation in Lau Basin, CO and methanol oxidation in Mid-Cayman Rise, and toluene and benzene degradation in Guaymas Basin (Fig. 1c, Fig. S7b). In parallel, the distribution and abundance of some microbial groups were also significantly enriched in the same samples accordingly (Fig. S7a) and underlay the functional differentiation, e.g., arsenate reduction in Lau Basin background deep-sea was attributed to members of Bacteroidetes and Thiomicrospirales while that same function in Lau Basin plumes was attributed to only Thiomicrospirales. CO oxidation in Mid-Cayman plumes was attributed to Chloroflexi, and toluene and benzene degradation in Guaymas Basin plume attributable to Methylococcales and Pseudomonadales (Supplementary Data 5). These observations are consistent with hydrothermal vent fluid geochemistry, e.g. Lau Basin hydrothermal vents have high arsenic end-member concentrations^33^ (ranging from 2.1-11 μmol/kg) and Guaymas Basin fluids contain aromatic hydrocarbons (primarily benzene and toluene)^34^.

As for within-site comparisons, the data indicate that the top three contributing taxa for major functions (including eight categories, carbon fixation, denitrification, sulfur cycling, hydrogen oxidation, methane oxidation, aerobic oxidation, iron oxidation, and manganese oxidation) are largely shared between plume and background deep seawater in Mid-Cayman Rise and Lau Basin, indicating functional consistency which was linked to taxonomy (Supplementary Data 5). Nevertheless, taxa abundance differed between plume and background, as reflected by both DNA and cDNA datasets associated with important functions (Supplementary Data 5, 6). Based on the results from energy landscape and MAG-based comparisons, our results suggest the adaptation of the plume microbiome, and demonstrate the consistency of links between taxonomy, function, and geochemistry.

### Sulfur cycling drives metabolic interactions in hydrothermal plumes

Building on our findings from both thermodynamic modeling and omics-based biogeochemical estimations which indicated the importance of sulfur-based metabolisms, we studied microbial metabolic interactions associated with sulfur cycling in all plumes. We recently developed a metric, metabolic weight score (MW-score)^35^ to measure the contribution of metabolic/biogeochemical steps, and their metabolic connectivity in a microbial community. More frequently shared functions and their higher abundances in a microbial community lead to higher MW-scores^35^. Both metagenomics and metatranscriptomic data showed elemental sulfur oxidation to be the key reaction in the sulfur cycle (Fig. 3a). In each community, sulfur oxidation had the highest MW-score (Fig. 4b, Fig. S10). Major contributors (*dsrAB* and *sdo* containing MAGs) to sulfur oxidation varied in different hydrothermal vent sites (Fig. 3b), indicating core sulfur oxidizers can have distinct distributions locally. Metabolic overlaps existed as some sulfur oxidizers had additional metabolic potential associated with utilizing various small carbon substrates and hydrogen, reducing nitrate/nitrite, and oxidizing iron/manganese/arsenite^36^ (Fig. 3c). Additionally, numerous connections of sulfur oxidation with other electron-transferring reactions were observed in the functional network (Fig. 4b c, d, and Fig. S10). Previously, sulfur-oxidizing bacteria belonging to SUP05 (Thiomicrospirales in GTDB R83 or PS1 in GTDB R202) and SAR324 lineages were identified to have metabolic plasticity involving the ability to conduct hydrogen oxidation and nitrate reduction^7, 37^ (in case of SUP05) and alkane/methane/carbon monoxide oxidation^17, 38^ (in case of SAR324) in plume and deep-sea environments, suggesting that plume microorganisms are optimized to mediate energy transformations upon available electron donors and acceptors. Here, our study indicates sulfur oxidizers are the primary group associated with energy scavenging from plume substrates. Sulfur oxidizers have metabolic plasticity to connect sulfur metabolism with other elemental transformations, are adapted to plume environments, and contribute significantly to biogeochemical cycles in the deep sea.

**Fig. 3.**
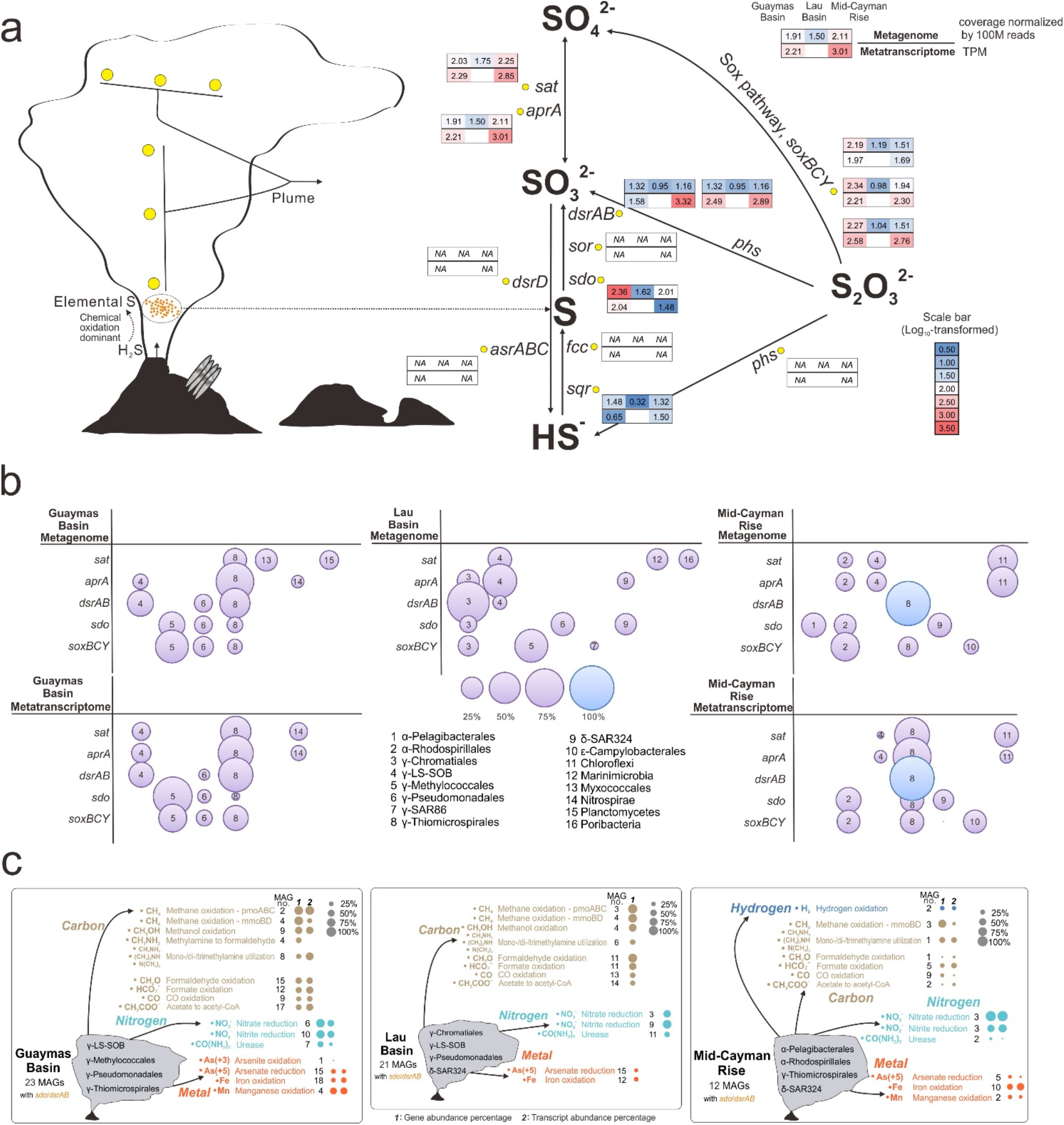
Sulfur metabolism and metabolic plasticity of sulfur oxidizers. **a** Details of sulfur metabolism pathways in the hydrothermal plume. The gene abundance (coverage normalized by 100M reads) and transcript expression level (TPM) for each step were calculated based on plume metagenomic and metatranscriptomic read mapping results. Log_10_ -transformed values of gene abundance and transcript expression level were labeled accordingly in the diagram. **b** Major contributors to sulfur metabolizing genes. For each sulfur metabolizing gene, microbial groups that occupied > 10% of the total gene abundance (by metagenome) or transcript expression (by metatranscriptome) values were labeled in the diagram. For some genes with only three or less than three contributors, all contributors were labeled. **c** Metabolic plasticity of sulfur oxidizers. For each hydrothermal vent site, three parameters were given to show the metabolic plasticity of sulfur oxidizers in conducting each electron transferring reaction related to carbon, nitrogen, hydrogen, and metal biogeochemical cyclings: the number of sulfur-oxidizing gene containing MAGs, gene abundance percentage, and transcript abundance percentage.

**Fig. 4.**
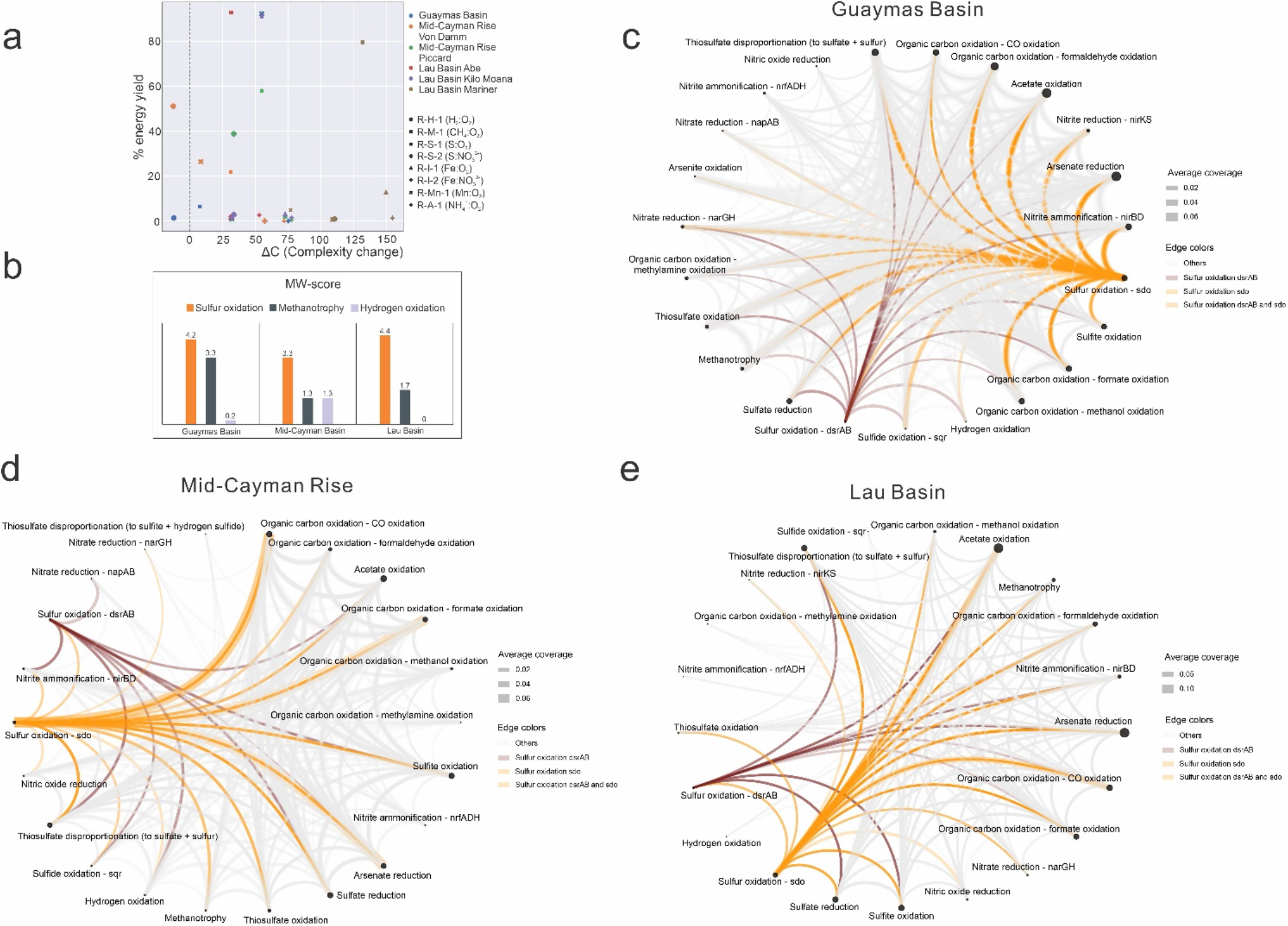
Network complexity, MW-scores (metabolic weight scores), and functional network diagrams of the three hydrothermal vent sites. **a** Network complexity diagram representing each reaction’s influence on the complexity of the network. In the figure, different colors represent different hydrothermal environments, different symbol shapes represent different reactions. The substrates (including electron donors and acceptors) were listed for each reaction in the legend. The x-axis is the change in complexity (ΔC) of the whole network for a node (a reaction here) and the y-axis is the percent energy yield of that reaction in the whole community. This network complexity diagram was based on thermodynamic estimation results at 3°C. **b** MW-scores of three major energy contributing reactions. **c** Functional network diagram of Guaymas Basin. **d** Functional network diagram of Mid-Cayman Rise. **e** Functional network diagram of Lau Basin. A group of metabolic cycling steps that are important in reflecting the plume substrate metabolisms were selected from METABOLIC-C regular MW-score results to make these functional network diagrams (**c, d, e**), respectively. In each functional network diagram, the size of a node is proportional to gene coverage associated with the metabolic/biogeochemical cycling step. The thickness of the edge represents the average gene coverage values of the two connected metabolic/biogeochemical cycling steps. Edges related to two reactions of sulfur oxidation were colored accordingly in each diagram.

While sulfur oxidation connects other metabolic reactions in the overall functional network and has significant energy yields, its role on the overall network complexity remains elusive. Specifically, we investigated the impact of sulfur metabolism on overall plume microbial metabolism. To address this, we built networks based on reactions and the percent energy yields, and investigated reaction influence on network complexity^39, 40, 41^ (Fig. 4a, Fig. S11). The network of reactions works as a whole mechanism where each reaction is one part^40^ and high ΔC reactions are key features of the networks. Most of these ΔC (complexity change) values are positive except for two points (Fig. 4a, Fig. S11). This indicates that all but two of these reaction nodes drive the system away from randomness and significantly contribute to the complexity of the network as a whole^40^. Meanwhile, in general, it seems that most reactions that are closer to smaller ΔC have higher percent energy yields associated with their reactions (Fig. 4a, Fig. S11). This phenomenon suggests that reaction nodes that result in higher changes of percent energy yields are not necessarily contributing to the reaction network’s complexity the most. Overall, this suggests that while sulfur oxidation tends to have higher energy yields, other reactions are also important components in plumes, and together cohesively contribute to the energy landscape.

### Low diversity, short migration history, and gene-specific sweeps in plume populations

Metagenomes provide full repertoires of genomic variation and facilitate interpreting fine-scale evolutionary mechanisms^42, 43, 44^. Here, we used *Tara* Ocean metagenomic datasets^45^ from the mesopelagic oceans to compare metagenomes from hydrothermal plume environments to the wider pelagic oceans and study the population genetic diversity of each MAG. We discovered that a large portion of MAGs exhibited a similar tendency of normalized single nucleotide variation (SNV) counts, nonsynonymous/synonymous substitution ratio of SNV (N/S SNV), and genome-wide mean *r*^2^ (Fig. 5a and Supplementary Data 11). In hydrothermal plumes, their SNV count is lower than *Tara* Ocean samples, N/S SNV ratio is higher than *Tara* Ocean samples, and mean *r*^2^ is higher than *Tara* Ocean samples. This suggests that in the plume: (1) Less SNVs are present, and population diversity is lower; (2) The population is younger with a short migration history. The higher N/S SNV ratio indicates that younger populations are less subjected to purifying (negative) selection to remove deleterious mutations; (3) The population is less subjected to recombination. The higher mean *r*^2^ reflects higher SNV linkage frequency at the genome-wide scale, indicating a lower recombination rate among population members.

**Fig. 5.**
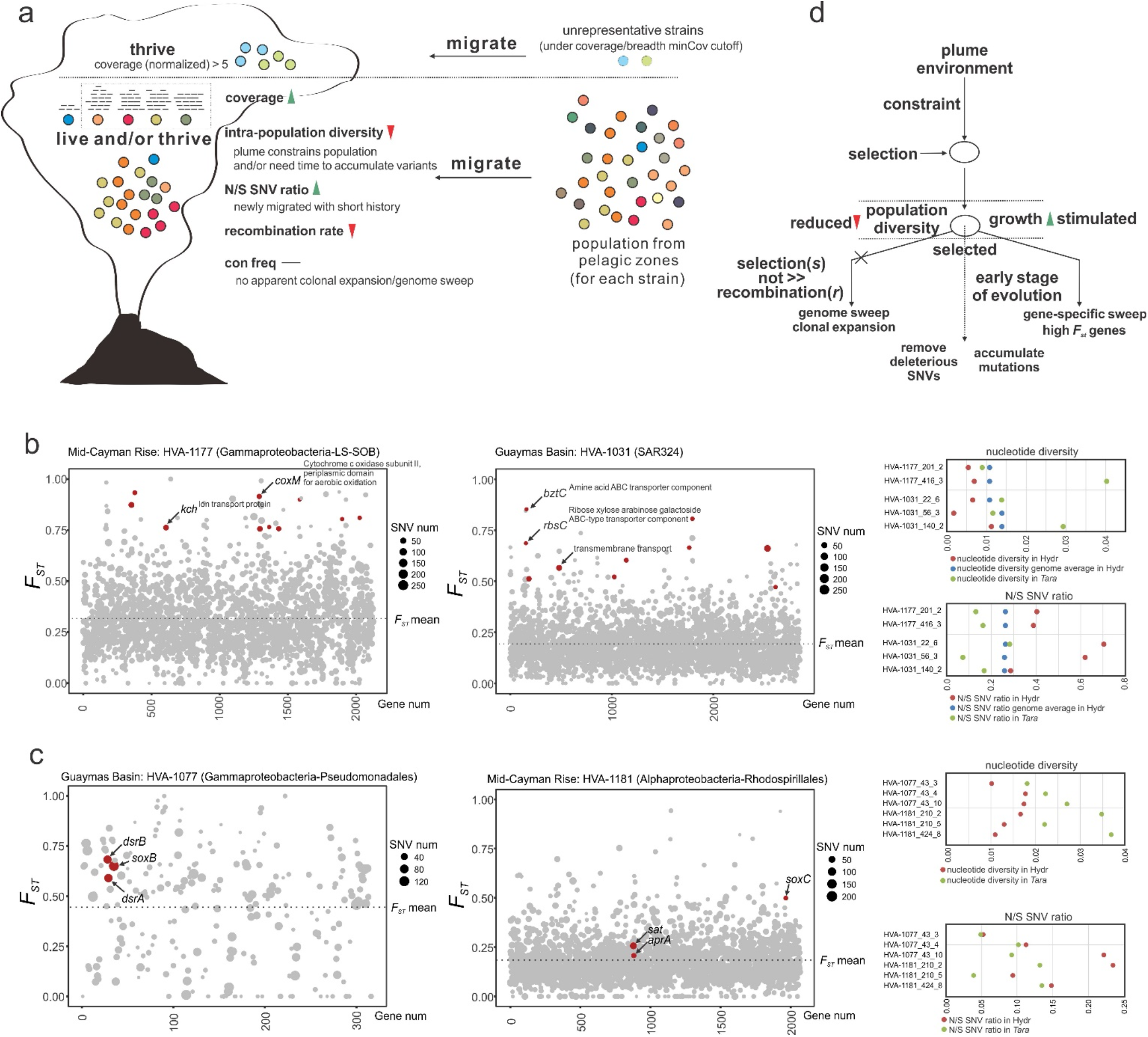
Evolutionary mechanism of plume microbial populations during migration. **a** Schematic diagram showing the changing trend of microdiversity parameters during migration. Individual solid dots with various colors represent microbial populations. Two scenarios were depicted in this panel: unrepresentative strains and strains that have detectable read mapping results in both environments. **b** Two representative charts showing *F*_*ST*_ distribution in MAGs that contain high *F*_*ST*_ genes. In each chart, the x-axis represents gene numbers (only genes with detectable *F*_*ST*_ ; negative values were removed). Dot sizes were proportional to SNV numbers in individual genes, and *F*_*ST*_ genome-wide mean was depicted in each chart with dash lines. Red-colored dots represent high *F*_*ST*_ genes that also passed the requirements of *F*_*ST*_, nucleotide diversity, N/S SNV ratios, and coverages (see methods). The nucleotide diversity and N/S SNV ratio distribution for high *F*_*ST*_ genes and genome-wide mean of all genes in different environments were depicted in the chart on the right side. Details of high *F*_*ST*_ genes and related parameters in individual genomes (all hits, also including these two representative genomes) were listed in Supplementary Data 12. **c** Two representative charts showing *F*_*ST*_ distribution in MAGs that contain sulfur metabolizing genes with signals of being fixed. In each chart, the x-axis represents gene numbers (only genes with detectable *F*_*ST*_ ; negative values were removed). Dot sizes were proportional to SNV numbers in individual genes, and *F*_*ST*_ genome-wide mean was depicted in each chart with dash lines. Red-colored dots represent sulfur metabolizing genes that passed the requirements of *F*_*ST*_, nucleotide diversity, N/S SNV ratios, and coverages (see methods). The nucleotide diversity and N/S SNV ratio distribution for sulfur metabolizing genes in different environments were depicted in the chart on the right side. Details of sulfur metabolizing genes with signals of being fixed and related parameters in individual genomes (all hits, also including these two representative genomes) were listed in Supplementary Data 13. **d** Frame diagram showing the underlying evolutionary processes during migration. Circles represent microbial populations. Dash line arrows indicate the direction of the next evolutionary step.

Next, we investigated potential signals of genome/gene sweeps using fine-scale evolutionary parameters. Consensus base frequency (frequency of reads supporting the consensus base), N SNV, and S SNV all showed no significant differences between plumes and the pelagic ocean (Supplementary Data 11). This indicates that these populations are unlikely to have undergone selective genome sweeps and clonal expansion during migration. We calculated fixation index *F*_*ST*_^46^ based on gene allele frequencies between these two environments (Fig. 5b and Supplementary Data 12) to investigate environmental selection. High *F*_*ST*_ genes are potential loci where selective pressures act on and indicate niche-specific adaptation. Further stringent criteria require the lower gene nucleotide diversity and higher N/S SNV ratio (Fig. 5b and Supplementary Data 12). Decreases of nucleotide diversity indicate gene-specific selective sweeps in the hydrothermal plume environment, and higher N to S SNV ratios suggest that these genes underwent a recent selection. Amongst 260 identified high *F*_*ST*_ genes using our stringent criteria, many of them involved transporters, aerobic oxidation, and stress responses (Fig. 5b and Supplementary Data 12). Transporters were associated with diverse substrates, e.g., metals (Co, Fe, and Mg), amino acids, Na^+^/H^+^, anions (nitrate/sulfonate/bicarbonate), carbohydrates (ribose/xylose/arabinose/galactoside), and aliphatic polyamines (spermidine/putrescine); meanwhile, these transporters are associated with many transporter families (Supplementary Data 12), including ABC superfamily, tripartite ATP-independent periplasmic (TRAP) family, tripartite tricarboxylate transporter (TTT) family, etc.

Given the observed importance of sulfur metabolism in plumes, we focused on the 238 identified sulfur metabolism genes. 23 of these genes had signals of being fixed after migration with *F*_*ST*_ values higher than the genome average (Fig. 5c and Supplementary Data 13). These genes were associated with sulfur oxidation, thiosulfate oxidation, and sulfite oxidation/sulfate reduction (*sat, aprA, sdo*, oxidative *dsrAB*, and *soxBC*) (Supplementary Data 13). This provides evidence that though not reaching the level of gene-specific selection sweeps, these genes were still being selected across the genome. Overall, this suggests a genetic adaptation to a sulfur-dominated environment after migration. An underlying evolutionary paradigm can be outlined from our population-level microdiversity analyses (Fig. 5c). As microbes enter the hydrothermal plume, some groups are selected for, and thrive due to substrates provided locally. This stimulates the growth of certain populations; meanwhile, constraints in the plume environment also induce selection effects and reduce the diversity of the population majority. Higher N/S SNV indicates they are young populations and are growing in the plume, consistent with the higher growth rates of major energy contributors. Gene-specific sweeps indicate local adaptation to the plume environment, and partially change population genetic structures after migration. Plume microbial populations are still in the early stage of evolution; as time goes on, mutations will progressively accumulate and deleterious SNVs will be gradually purged.

## Discussion

In this study, we observed that distinctive plume geochemistry influences the energy landscape across three different hydrothermal vent sites^4, 29^. Sulfur oxidation is the major energy-yielding reaction, while different sites are also represented by different energy landscapes influenced by differing vent geochemistry. For instance, other important energy sources like methane and hydrogen also have important roles in the energy landscape of hydrothermal plumes. The existence of a core plume microbiome indicates that a general biogeochemical feature – energy and substrate supply – within hydrothermal plumes supports the growth of these globally dispersed cosmopolitan microorganisms. As a consequence, the core plume microbiome is likely a result of the sulfur oxidation-based energy landscapes shared among many hydrothermal plumes around the globe. The increased taxa abundance and higher growth rates of major energy contributing taxa supports the interpretation that microbiomes act in response to geochemically-influenced energy landscapes with some taxa fueled by plume substrates. These analyses support the theory of an ocean seed bank origin of the hydrothermal plume microbiome^47^. Plume geochemistry defines the energy availability, serving as a key control on the microbiome distribution and abundance^2, 9^. The consistent taxonomy-function-geochemistry links demonstrated by us suggest that omics-based profiling that reflects the full genetic repertoire of plume microorganisms can be a powerful tool to unravel the relationship between environments and microbiomes.

Characterization of sulfur metabolism in plumes reveals that though all plumes have sulfur oxidation as the reaction with the highest MW-score, and sulfur-oxidizing genes were highly expressed, the major populations contributing to these processes (*dsrAB* and *sdo* containing MAGs) vary in different hydrothermal vent sites. This indicates the variable composition of core sulfur oxidizers in individual environments which suggests the endemicity of microbial community structure. Core sulfur oxidizers can be derived from the pelagic ocean through stochastic processes that can be influenced by dormancy capacity to provide resilient seed microbes, ocean currents to overcome dispersal limitations, and adaptive strategies to nutrient and temperature fluctuations^2^. Core members of the plume microbiome derived in this manner likely thrive under favorable geochemical conditions^48^. For example, Pseudomonadales, Thiomicrospirales, and SAR324 are members of the core plume microbiome, but are also known to be abundant cosmopolitan bacteria in the pelagic oceans. These microorganisms can be distributed as seed banks in the global oceans, triggered by plume sulfur substrates, and subsequently become active sulfur oxidizers in hydrothermal plumes^9, 48^. Sulfur oxidizers within the community have metabolic plasticity to connect other energy transformation activities, e.g., small carbon substrate utilization, nitrate/nitrite reduction, and iron/manganese/arsenite oxidation, etc. This indicates that sulfur and other energy sources can be simultaneously utilized for energy conservation by sulfur oxidizers even in various plume environments with different energy landscapes. At the same time, as described in the network complexity analysis, though sulfur oxidation dominates in energy generation, other reactions are also important components in the metabolic network connected to sulfur oxidation, and cohesively contribute to the energy landscape.

Finally, the microdiversity patterns observed in plume microorganisms depict a scheme of populations selected by environmental constraints. Low population diversity and high N/S SNV ratio indicate that microbes are selected by plume conditions and actively grow after a short migration history. Evidence shows that gene-specific sweeps within certain plume populations are related to nutrient uptake, aerobic oxidation, and stress responses, and some sulfur metabolizing genes are also selected during the environmental change. These traits help microbial cells to be more adaptable and resilient in sulfur oxidation-dominated hydrothermal plume conditions. Collectively, the plume microbiome has a distinctive composition, function, and genetic structure focused on allowing organisms to better adapt to hydrothermal plume conditions. Population alteration in plumes compared to the background deep sea involves both reshaping community-level structure and fine-scale strain-level genetic adjustments that includes advantageous metabolisms being fixed. These nuanced microdiversity changes can lead to fundamental changes in population fitness towards niche adaptation. Overall, the plume microbial community is associated with energy conservation, metabolic distribution, and cell stress response which likely facilitates more efficient adaptation of the plume microbiome in mediating biogeochemical cycles. The connected relationship between microbiome and biogeochemistry we demonstrate reflects the overall ecological and evolutionary basis of microbial strategies for thriving in geochemically-rich energy landscapes.

## Data availability

The MAG genomic sequences are deposited into the NCBI Genome database under the BioProject ID of PRJNA488180. The genome annotation results from this study are publicly available at https://doi.org/10.5281/zenodo.5034800 (all plume MAG annotations are deposited to this location).

## Code availability

The Perl and R codes for parsing, calculating, and visualizing in this study are publicly available at https://github.com/AnantharamanLab/Hydrothermal_plume_omics_Zhou_et_al._2021.

## Acknowledgments

We thank all the members in the cruise of R/V New Horizon in Guaymas Basin, Gulf of California (July 2004), R/V Atlantis and R/V Falkor in Mid-Cayman Rise, Caribbean Sea (Jan 2012 and June 2013), R/V Thomas G Thompson in Eastern Lau Spreading Center (ELSC), Lau Basin, western Pacific Ocean (May-July 2009), R/V MGS Sagar in Central Indian Ridge and Southwest Indian Ridge (Jan-Mar 2017), and R/V Thomas G Thompson in Axial Seamount (Aug 2015) in assisting sampling and physicochemical data measuring and processing. This research was supported by the National Science Foundation under grant number OCE 2049478 to K.A and OCE 1851208 to J.A.B and the Gordon and Betty Moore Foundation under grant number GBMF3297 to J.A.H.

## Author contributions

Z.Z. and K.A. conceived the project. K.A. designed the framework of this project. Z.Z. led the research and conducted the data analysis. Z.Z. and K.A. wrote the manuscript. J.A.B. and G.J.D. sampled in the cruises and J.A.B. contributed to the thermodynamic modeling. K.K., P.Q.T, and A.M.A. helped on data analysis and/or visualization. All authors (Z.Z., P.Q.T., A.M.A., K.K., J.A.B., R.K.S., K.P.K., P.J.K., C.S.F., C.S.S., J.A.H., M.L., G.J.D., and K.A.) reviewed the results, revised, and approved the manuscript.

## Additional information

Supplementary Information is available for this paper on XXX website.

**Reprints and permissions information** is available at www.nature.com/reprints

**Correspondence and requests for materials** should be addressed to K.A.

## Methods

### Sample information and omics sequencing

Hydrothermal plume and surrounding background samples were collected from the corresponding cruises: R/V New Horizon sampling in Guaymas Basin, Gulf of California (July 2004), R/V Atlantis and R/V Falkor sampling in Mid-Cayman Rise, Caribbean Sea (Jan 2012 and June 2013), two consecutive cruises on the R/V Thomas G Thompson sampling in Eastern Lau Spreading Center (ELSC), Lau Basin, western Pacific Ocean (May-July 2009), R/V MGS Sagar sampling in Central Indian Ridge and Southwest Indian Ridge (Jan-Mar 2017), and R/V Thomas G Thompson sampling in Axial Seamount, Juan de Fuca Ridge, northeastern Pacific Ocean (Aug 2015). In brief, Guaymas Basin plume and background samples were collected by 10 L CTD-Rosette bottles and N_2_ -pressure filtered on board for microbial specimen collection by 0.2 µm pore size, 142 mm polycarbonate membranes^11^. The samples were preserved immediately in RNAlater. Mid-Cayman hydrothermal plume and surrounding background samples were collected by Suspended Particulate Rosette (SUPR) filtration device^49^ mounted to the remotely operated vehicle Jason II. SUPR collected water with the amount of 10-60 L from different sampling locations, and these samples were *in-situ* filtered for microbial specimens by 0.2 μm pore size SUPOR polyethersulfone membranes and preserved in RNAlater flooded conical vials and frozen at -80°C. For Lau Basin samples, SUPR-collected samples were *in-situ* filtered by SUPOR polyethersulfone membranes with 0.8 μm and 0.2/0.8 μm pore size for geochemical analysis and microbial specimen collection, respectively^26^. Samples were preserved in RNAlater flooded conical vials and frozen at -80°C. Central Indian Ridge and Southwest Indian Ridge samples were collected by 10 L CTD-Rosette bottles and filtered by 0.2 µm pore size, 47 mm SUPOR polyethersulfone membranes, and preserved in RNAlater flooded conical vials and frozen at -80°C. For Axial Seamount samples, both plume and background samples were collected by a Seabird SBE911 CTD and 10L Niskin bottles^50^. Samples of 3 L were then transferred into cubitainers, filtered through 0.22 μm Sterivex filters, and preserved for downstream analysis^50^.

Details for sample collection, preservation, geochemical analysis, and metagenomic/metatranscriptomic sequencing refer to the previous publications^22, 50, 51^. Detailed cruises and sampling information refer to Supplementary Data 1. The geological map and schematic diagram represent the details of sampling locations (Fig. 1a, Fig. S1). The metagenomic DNA and metatranscriptomic cDNA were extracted and synthesized from corresponding samples and processed for Illumina HiSeq 2000/2500 sequencing as described previously^11, 14, 18, 25, 50^. The distribution of acquired metagenomes (DNAs, labeled as “D”) and metatranscriptomes (cDNA, labeled as “C”) was represented in Fig. S1b (only for samples with detailed location and physicochemical characterization; distribution of other samples refers to Supplementary Data 1). The raw reads (both DNA/cDNA reads) were dereplicated by SeqTools v4.28 (https://www.sanger.ac.uk/tool/seqtools/) and processed by Sickle v1.33 (https://github.com/najoshi/sickle) to trim reads of low quality with default settings. Command “reformat.sh” in BBTools (last modified on Feb 11, 2019; https://www.sourceforge.net/projects/bbmap/) was used to calculate fastq sequence and nucleotide numbers.

### Core hydrothermal plume microbiome analysis

In total, 51 hydrothermal plume and background 16S rRNA gene datasets were used for analyzing the microbiome of hydrothermal plume, within which 24 datasets were obtained in this study, containing datasets from samples of Mid-Cayman Rise, Guaymas Basin, Lau Basin, Central and Southwestern Indian Ridge, and Axial Seamount plume (Supplementary Data 2). For hydrothermal plume and background samples with only metagenome datasets, 16S rRNA gene sequences were parsed out from metagenomes and these sequences were weighted according to their coverages. Simulated 16S rRNA gene datasets were used in subsequent analysis. The original datasets of paired-end reads were merged into combined 16S tags by FLASH v1.2.11^52^ with default settings. The bioinformatic analyses, including pre-analysis quality control, 16S chimera checking, open-reference OTU picking, taxonomy assigning, OTU table file ‘biom’ generating and rarefying, OTU representative sequences filtering and aligning, alignment filtering, and phylogenetic tree reconstructing, were performed according to the instructions of QIIME v1.9.1^53^, respectively. The 16S rRNA reference database was based on “SILVA_132_QIIME_release”^54^. The resulted ‘biom’ (OTU table file), ‘tre’ (phylogenetic tree), and “map” (sample characterization map) files were imported into R (using R package ‘*phyloseq*’) for downstream analysis and visualization. Taxa summary and principal coordinates analysis (PCoA) were conducted accordingly to delineate the community structure and biogeographic pattern of hydrothermal plume and background seawater microbiome. Genus-level taxa summary table was used to find core hydrothermal plume microbiome from 37 hydrothermal plume datasets by filtering genera that exist in > 67% plume datasets and have > 1% relative abundance on average. Core plume microbiome metabolic profiles were conducted by choosing MAGs (see the following sections of obtaining these MAGs) from this study that contain 16S rRNA genes affiliated to the core plume microbial genera. Metabolic profiling for these MAGs was based on the result from “MAGs genomic property and annotation”.

### Assembling and metagenomic binning

QC-processed reads were assembled *de novo* by MEGAHIT v1.1.2^55^ with settings as “--k-min 45 --k-max 95 --k-step 10”. Hydrothermal plume and background metagenomes from the same hydrothermal site were assembled together. QC-processed reads were re-mapped to assemblies by Bowtie 2 v2.2.8^56^ with default settings. For each hydrothermal site, hydrothermal plume and background reads were mapped to corresponding assemblies separately; bam files by plume and background samples for individual assemblies were used for downstream binning, Subsequently, the assemblies were subjected to a MetaBAT v0.32.4^57^ based binning with 12 combinations of parameters. Afterward, DAS Tool v1.0^58^ was applied to screen MetaBAT MAGs, resulting in high quality and completeness MAGs. This MetaBAT/DAS Tool method enables a comprehensive “slice-layer profiling” for searching potential MAGs with a better outcome (in-house tested). CheckM v1.0.7^59^ was used to assess MAG quality and phylogeny. Outlier scaffolds with abnormal coverage, tetranucleotide signals, and GC pattern within potential high contamination MAGs (by CheckM) and erroneous SSU sequences within MAGs were screened out and decontaminated by RefineM v0.0.20^60^ with default settings. Afterwards, further MAG refinement for decontaminating certain MAGs was manually inspected based on VizBin^61^. MAGs are picked using a threshold of < 10% contamination (namely genome redundancy) and > 50% completeness.

### MAG genomic property and annotation

Genome phylogeny was determined by RefineM and GTDB-Tk v 0.2.1^62^ (GTDB database, release 83). Additionally, phylogenies of those genomes that could not be assigned to a meaningful microbial group were inferred from ribosomal protein (RP) trees using the phylogenetic reconstruction method described below. Genomic properties, including genome coverage, genome and 16S rRNA taxonomy, tRNAs, genome completeness, and scaffold parameters, were parsed from results that were calculated by CheckM and tRNAscan-SE 2.0^63^. Relative genome coverages were normalized by setting each metagenomic dataset size as 100M paired-end reads. MAG ORFs were parsed out by the Prokka annotation pipeline v1.12^64^ with default settings. For ORF annotation, GhostKOALA v2.0^27^, KAAS v2.1^26,^ and eggNOG-mapper v4.5.1^28^ were applied to thoroughly annotate ORFs to KOs. For eggNOG-mapper annotation, we used its first KO hit as annotation result; if there was only COG annotation, we translated it into KO using ‘ko2cog.xl’ provided by KEGG database. When combining three software annotations, we use resulted KO from the first software as the final annotation; if there is no annotation from the first software, then we will move to the next software accordingly. Annotation by NCBI nr database (Mar 6, 2017 updated) was conducted with default settings and for each annotation the first meaningful hit (hit not assigned as ‘hypothetical protein’) was extracted. Genomic-specific metabolic traits were searched against TIGRfam, Pfam, Kofam, and custom HMM profiles using hmmscan^65^ and custom protein database using DIAMOND BLASTP^66^. For searching against custom HMM databases, noise cutoff values are determined according to previous settings^12^, respectively. For DIAMOND BLASTP searching, a stringent criterion as “-e 1e-20 --query-cover 65 --id 65” was applied. Carbohydrate active enzymes (CAZymes) were searched against dbCAN2 with default settings^67^; Peptidases were searched against MEROPS ‘pepunit’ database with stringent DIAMOND BLASTP settings as “-e 1e-10 --subject-cover 80 --id 50”^68^.

### Phylogenetic tree reconstruction

The syntenic block of universal 16 ribosomal proteins (RPs) (L2-L6, L14-L16, L18, L22, L24, S3, S8, S10, S17, and S19) were used for inferring RP phylogenetic tree, after hmmscan-based^65^ searching for RPs from all MAGs. The individual RP was pre-aligned with local custom RP database by MAFFT v7.123b^69^ and curated in Geneious Prime v2019.0.4^70^ by manually masking out begin and end regions with lots of gaps. Out of 206 MAGs, 177 containing > 4 RPs were used; the concatenated and curated 16RP-alignment (7741 aligned columns) was used for phylogenetic inference by IQTREE-based maximum likelihood method (IQ-TREE multicore v1.6.3^71^) with settings of “-m MFP -bb 1000 –redo -mset WAG,LG,JTT,Dayhoff -mrate E,I,G,I+G -mfreq FU -wbtl”. The resulted phylogenetic tree was rooted by archaea lineages and visualized by iTOL^72^. Functional traits were added accordingly to each MAG on the tree. Bacterial and archaeal SSU sequences (> 300 bp and the longest from individual MAG) parsed out by local pipeline (use CheckM ssuFinder^59^ to pick and RefineM to filter erroneous hits) were aligned in SINA aligner^73^ with default settings. The 16S sequence taxonomy was checked by BLASTn searching against SILVA_128_SSUParc_tax_silva database^54^ and 16S sequences with resulted taxonomy different from their MAG phylogeny (at the phylum level) were filtered due to the high possibility of contamination. IQTREE-based^71^ phylogenetic inference was conducted with settings of “-st DNA -m MFP -bb 1000 -alrt 1000”. The 16S rRNA gene tree based on the alignment of 85 sequences with 50000 columns was rooted by archaea lineages, visualized by iTOL^72^, and manually curated.

### Metagenomic and metatranscriptomic mapping

QC-passed metagenomic reads were mapped to MAGs separately (metagenomic datasets from Guaymas Basin, Mid-Cayman Rise, and Lau Basin sites were mapped individually to the corresponding MAGs) using Bowtie 2 v2.2.8 with default settings^56^. MetaBAT integrated “jgi_summarize_bam_contig_depths” script and homemade Perl scripts were used to calculate MAG coverage (normalized coverage with each metagenomic dataset size set as 100M paired-end reads). QC-passed metatranscriptomic reads (use the same QC-process as described above with an additional SortMeRNA v2.1^74^ rRNA filtering step) were mapped to MAGs separately, with TPM (Transcripts Per Kilobase Million) calculated for individual genes within each genome.

### Statistical comparison on MAG and functional trait abundance

Metagenome/metatranscriptome-based MAG mapping results and functional annotations for all the MAGs were summarized individually. Afterwards, significance tests on the differentiation pattern of MAG (also MAG taxonomic group) and functional trait abundances across all the metagenomic/metatranscriptomic samples were calculated by R package DESeq2^75^. Log2 Fold Change value with adjusted *P*-value (by nbinomWaldTest) < 0.05 was considered as significant. Relative abundances of MAG (also MAG taxonomic group) and functional traits were visualized by R (using R package ‘*pheatmap*’) with the relative abundance at row normalized by removing the mean (centering) and dividing by the standard deviation (scaling). Sunburst figures were generated to depict the relative abundance of MAGs based on metagenomic/metatranscriptomic mapping results, with the significant Log2 Fold Change values labeled to individual MAGs that have differential abundances between different hydrothermal ecological niches, e.g., plume and background.

To find taxa in microbial community that are responsible for enriched functions (functions that are significantly enriched in each environment), major functions (including functions that are in the categories of carbon fixation, denitrification, sulfur cycling, hydrogen oxidation, methane oxidation, aerobic oxidation, iron oxidation, and manganese oxidation), and specific functions, custom Perl scripts were written to get the corresponding microbial community contribution information (scripts deposited in https://github.com/AnantharamanLab/Hydrothermal_plume_omics_Zhou_et_al._2021). Functional trait results of all MAGs, MAG coverage within the community, and targeted function list were used as inputs to conduct the calculation. For environments with metatranscriptomic reads, we also used active MAG coverage (calculated by metatranscriptomic reads mapping result) as the input to calculate microbial community contribution information based on metatranscriptomes.

### Bioenergetic and thermodynamic modeling

Equilibrium thermodynamic reaction path modeling was used to predict chemical concentrations and activity coefficients resulting from the mixing of seawater with end-member vent fluids (Supplementary Table 2). Our thermodynamic modeling builds on the specific plume model implementation described in Breier et al^76^. The estimated temperature of bottom seawater is according to the previous report^10^. The original chemical data is derived from Reeves et al^77^ and Anantharaman et al^10^. For each hydrothermal vent system, we choose at least one representative end-member fluid sample(s), respectively (1 for Guaymas Basin, 2 for Mid-Cayman Rise, and 3 for Lau Basin) (Supplementary Table 2).

Bioenergetic and thermodynamic modeling procedures were conducted as described in Anantharaman et al^7^ and Li et al^18^ (More details refer to Supplementary Information and Tables). Reaction path modeling was performed with REACT, part of the Geochemist’s Workbench package^78^. Conductive cooling was neglected and mixture temperatures were a strict function of conservative end-member mixing. Precipitated minerals were allowed to dissolve and their constituents to re-precipitate based on thermodynamic equilibrium constraints. Thermodynamic data were predicted by SUPCRT95^79^ for the temperature range of 2°C to end-member vent fluid temperature and a pressure of 500 bar. The estimated biomasses and free energies of individual environments were calculated and their relative abundance change along the temperature range (2 - 121°C) was visualized by R. Two temperatures (3 and 4.9°C) were picked to conduct the biomass and free energy estimation for representing typical plume temperatures in nature.

### Energy contribution and MAG growth rate calculation

Based on metabolic prediction of each MAG and MAG gene coverage and expression level within each environment, energy contribution for each electron donor was calculated based on gene coverage/expression level and free energy of each catabolic reaction. The contribution ratio of electron donor species was calculated for individual samples respectively. We also included influence of the presence of electron acceptors to energy contribution calculation. To simplify the hydrothermal condition, we only included two major electron acceptors (O_2_ and NO_3_^-^) and used the ratio of these two electron acceptors to infer energy contribution of electron donors at different oxidative conditions.

Microbial genome replication starts directionally from a single origin^31^. Based on metagenomic mapping, at a single time-point the coverage ratio between the replicating origin and terminus of a microbial genome can be used as a proxy to represent the replication rate/growth rate^30, 32^. Growth rate for each MAG was calculated by iRep v1.10^30^ with default settings. MAGs that are from the same environments were pooled together as the input genomes. Sam files that were generated by metagenomic mapping described above were used as the iRep input. Barcharts that reflect the growth rate and significant difference test result (by *t*-test) of MAG taxonomic groups were generated using R package ‘*ggplot2*’ and ‘*PairedData*’.

### Network complexity analysis

For each community, a bipartite network was built based on reaction/substrate relationships and the percent energy yields for each reaction. Briefly, the plume chemical reaction table for each reaction was stored; within the table, the substrate and product for a reaction were recorded^39^. Then, for each community, reactions (represented as one set of nodes in the bipartite network) with different percent energy yields were connected with substrates and products in the network (represented as the second set of nodes) via directed edges between both sets of nodes. The energy yields are based on the result from “Bioenergetic and thermodynamic modeling” and are represented on the network as node size proportional to the percent energy yield. These networks were constructed using the Python package ‘*networkx*’^80^ (https://networkx.org/).

The network complexity change as a function of reaction energy yield was calculated by the following steps^40^. For each plume community network, the complexity of the network’s structure was measured. A node was taken from the network; as a consequence, the change in complexity (ΔC) before and after the node was taken was calculated accordingly. The ΔC was assigned to that node as a property representing that node’s contribution to the network’s overall complexity. Then this node was placed back and these steps were repeated for each reaction node^40^.

In this study, complexity (C) was calculated by estimating the algorithmic complexity. Because algorithmic complexity cannot be directly computed, we used an estimate known as the <u>B</u>lock <u>D</u>ecomposition <u>M</u>ethod (BDM)^41^. The perturbation analysis to calculate each node’s complexity contribution (ΔC) is called <u>M</u>inimal <u>I</u>nformation <u>L</u>oss <u>S</u>election, MILS^32^; in this study, successive edge deletion was replaced as node deletion which also works with good performance^33^. This method has been used to characterize complex properties of biological networks and is proven to be a good measure among many other algorithms^40, 41^. For all reaction nodes in each community plume reaction network, we conducted this measurement for each reaction node and came up with the scatterplots.

### Community-level metabolic analysis

Resulted MAGs and plume metagenomic reads were used to conduct community-level metabolic analysis using METABOLIC-C v4.0^35^ with default settings. For Guaymas Basin, Mid-Cayman Rise, and Lau Basin sites, all MAGs and plume metagenomic reads from each site were used separately. From METABOLIC-C regular MW-score results, a group of metabolic cycling steps that are important in reflecting the plume substrate metabolisms were specifically selected to make functional network diagrams (using R script ‘*draw_functional_network*.*R’* from METABOLIC-C). For each site, MW-score table and functional network diagram (based on both all and selected metabolic steps) were generated, respectively.

### Evolution analysis

Metagenomic reads from mesopelagic *Tara* Ocean metagenomic datasets (with > 800m depth)^45^ were used as the regular ocean environment representatives to compare microdiversity characteristics with that of hydrothermal environments from this study. To simplify analyses, *Tara* Ocean reads from samples collected by filtration with various filter sizes at each station were pooled as one to represent all reads from that station. Both *Tara* Ocean reads and hydrothermal environment reads (including both background and plume environments; background and plume reads were also pooled together individually to simplify analyses and satisfy coverage requirement of each MAG) from this study were first mapped to hydrothermal environment MAGs recovered from individual sites by Bowtie 2^56^ with default settings. After mapping, reads within resulted bam files were filtered according to the following rules to calculate downstream microdiversity parameters: (1) minimum percent identity of read pairs to reference > 95%; (2) maximum insert size between two reads < 3× median insert size and minimum insert size > 50bp (so only paired reads are retained). Filter steps were either conducted by inStrain v1.4.1^42^ or inStrain_lite v0.4.0^81^ (for generating bam files) with the same rules. Software inStrain was further employed to calculate microdiversity parameters for each MAG in individual sites from this study. Subsequently, interested parameters^42^ were picked and parsed accordingly from resulted folders, including ‘coverage’ (average coverage depth of all scaffolds of one genome), ‘breadth minCov’ (percentage of bases in the scaffold that have at least ‘min_cov’ coverage), ‘SNV count / (breadth minCov × length)’ (total number of SNVs called on one genome normalized by genome length and breadth minCov), ‘N/S SNV ratio’ (nonsynonymous to synonymous SNV ratio of one genome), ‘r2_mean’ (*r*^2^ mean between linked SNVs), ‘con freq mean’ (mean value of fraction of reads supporting the consensus base within one genome), ‘con freq mean for N SNV’ (mean value of con freq on all nonsynonymous SNV sites), and ‘con freq mean for S SNV’ (mean value of con freq on all synonymous SNV sites). MAGs that have breadth_minCov value < 50% or do not pass the ‘min_cov’ requirement by inStrain were removed from microdiversity analysis in each site.

In order to identify gene-specific selective sweep in hydrothermal environment, we further pooled reads together into two categories, one contains hydrothermal environment datasets (including both background and plume environment datasets) and the other contains *Tara* Ocean samples (all *Tara* Ocean sample datasets are pooled together). After reads mapping and filtering as described above, *F*_*ST*_ (fixation index) between hydrothermal and *Tara* Ocean environments was calculated using scikit-allel package^82^ (Hudson method^83^) within inStrain_lite to identify genes with skewed allele frequencies across the whole genome. Subsequently, high *F*_*ST*_ genes from each MAG within each hydrothermal vent site were identified if they have *F*_*ST*_ value > *F*_*ST*_ mean (genome-wide *F*_*ST*_ average) + 2.5 × *F*_*ST*_ std (genome-wide *F*_*ST*_ standard deviation) and the lowest gene coverage in either hydrothermal and *Tara* Ocean environment samples should be higher than 5×. Meanwhile, for each genome the number of genes with empty *F*_ST_ value should not be more than half of all genes, otherwise high *F*_*ST*_ genes will not be taken into account for this genome. We set gene coverage in both environments to be at least 5× due to the fact that reduction of gene coverage (or loss of coverage in some genome regions) can also lead to low nucleotide diversity. Furthermore, to confirm that these genes are specifically selected in hydrothermal environment, additional requirements were added: (1) gene nucleotide diversity in hydrothermal environment < nucleotide diversity genome average in hydrothermal environment; (2) gene N/S SNV ratio in hydrothermal environment > N/S SNV ratio genome average in hydrothermal environment; (3) gene nucleotide diversity in hydrothermal environment < gene nucleotide diversity in *Tara* Ocean samples; (4) gene N/S SNV ratio in hydrothermal environment > gene N/S SNV ratio in *Tara* Ocean samples.

To find sulfur metabolizing genes that have signals of being fixed after migration, a relatively less stringent set of criteria were used to screen gene *F*_*ST*_ values compared to high *F*_*ST*_ gene identification method in the above paragraph. For each sulfur metabolizing gene (including genes of *sat, aprA, sdo*, oxidative *dsrAB*, and *soxBCY*) containing MAGs, the identified genes should meet the following criteria: (1) *F*_*ST*_ value > *F*_*ST*_ mean (genome-wide *F*_*ST*_ average) and both *F*_*ST*_ and *F*_*ST*_ mean should be positive values; (2) gene nucleotide diversity in hydrothermal environment < gene nucleotide diversity in *Tara* Ocean samples; (3) gene N/S SNV ratio in hydrothermal environment > gene N/S SNV ratio in *Tara* Ocean samples; (4) gene coverages in hydrothermal environments and *Tara* Ocean samples both > 5×. Sulfur metabolizing genes that meet all the four criteria were indicated to have positive gene fixation signals though the selective power across the genome did not reach the level of gene-specific selective sweeps as indicated by the above method.

