## Supplementary Information and Tables 1-3 for "The sulfur cycle connects microbiomes and biogeochemistry in deep-sea hydrothermal plumes"

### Methods

#### Bioenergetic and thermodynamic modeling

To predict the chemical concentrations and activity coefficients in the plume, the equilibrium thermodynamic reaction path modeling approach was adopted. The modeling processes were imposed on the mixtures of seawater and end-member vent fluids from individual hydrothermal environments, including Guaymas Basin, Lau Basin, and Mid-Cayman Rise. The chemical parameters for these seawater and vent fluids samples are provided in Supplementary Table 2. Our thermodynamic modeling builds on the specific plume model implementation of Breier et al^1^. The estimated temperature of bottom seawater was adopted according to the previous reports^2^. The original chemical data was derived from Reeves et al^3^. For each hydrothermal vent system, we choose at least one representative end-member fluid sample(s), respectively (1 for Guaymas Basin, 2 for Mid-Cayman Rise, and 3 for Lau Basin) (Supplementary Table 2). The following is a brief description of how the modeling was conducted in this study, as modified from the detailed description in Anantharaman et al^4^.

The equilibrium thermodynamic reaction path modeling was based on the mixing process with the ratio of vent fluids vs. seawater as 1: 10,000. This ratio was chosen to resemble the dilution ratio of vent fluids at the non-buoyant plumes in this study. The reaction path modeling was conducted by REACT, a constitutional program implemented in the Geochemist’s Workbench package^5^. The thermodynamic prediction (both the Gibbs free energy and biomass yields) was calculated by SUPCRT95^6^ with the temperature range of 1–425°C and the pressure of 500 bar which can cover all known deep-sea hydrothermal vents. The available energy per kilogram plume fluid was estimated by calculating ΔG for the metabolic reactions listed in Supplementary Table 3, using the results of the reaction path model and multiplying ΔG by the concentration of the most limiting reactant. Resulting Gibbs free energy and biomass yields are reported on a per kilogram plume fluid basis.

**Supplementary Table 1. Membership of the core plume microbiome.** The metabolic characteristics (including capability based on genomes or distribution patterns) were based on currently available reports/publications of the same or sub-microbial groups.

| **Microbial group** | **Phylogeny** | **Metabolic characteristics** | **Potential origin** |
| --- | --- | --- | --- |
| Sva0996 marine group | Actinobacteriota | uncultured marine microorganisms | likely seawater^7^ |
| *Sulfurimonas* | Epsilonbacteria; Campylobacterales | reduce nitrate, oxidize both sulfur and hydrogen^8^ | seafloor sediment/subsurface^9^ |
| SAR202 clade | Chloroflexi | specifically inhabit the aphotic realm; members metabolize multiple organosulfur compounds; many appear to be sulfite-oxidizers^10^ | seawater^10^ |
| Marinimicrobia | Marinimicrobia | members contain N_2_O reductase, nitrate reductase, and polysulfide reductase^11^ | seawater^11^ |
| JL-ETNP-F27 | Planctomycetota | uncultured marine microorganisms | likely seawater^12^ |
| Pla3 lineage | Planctomycetota | uncultured marine microorganisms | likely seawater |
| Magnetospiraceae | Alphaproteobacteria; Rhodospirillales | magenetotactic, chemoorganoheterotrophic and chemolithoautotrophic under microaerobic condition (based on *Magnetospira thiophila*)^13^ | likely seawater^13^ |
| *Alteromonas* | Gammaproteobacteria; Alteromonadales | chemoorganotrophic, aerobic^14^ | seawater^14^ |
| *Marinobacter* | Gammaproteobacteria; Alteromonadales | utilize a variety of aliphatic and aromatic compounds, both aerobic or anaerobic with nitrate/nitrite^14^ | seawater^14^ |
| *Pseudomonas* | Gammaproteobacteria; Pseudomonadales | capable of heterotrophic Mn(II)-oxidation^15^; members from hydrothermal sediment are potential PAH degraders^16^ | seawater or marine sediment |
| HOC36 | Gammaproteobacteria | sponge/coral symbiotic microorganisms^17^^,^ ^18^ | seawater^26^^,^ ^27^ |
| SAR86 clade | Gammaproteobacteria | chemoheterotrophic, aerobic; capable of degrading lipids and polysaccharides^19^; conducting proteorhodopsin-based photosynthesis^19^ | seawater^19^ |
| SUP05 cluster | Gammaproteobacteria; Gammaproteobacteria *incertae sedis* | deep-sea hydrothermal SUP05 cluster can oxidize sulfur and hydrogen^4^ | seawater^4^ |
| SAR324 clade | Deltaproteobacteria | degrade aliphatic and aromatic hydrocarbon, and alcohol; oxidize sulfur, methane, and formate^20^^,^ ^21^ | seawater^20^^,^ ^21^ |

**Supplementary Table 2. Chemical parameters of hydrothermal vent end-member fluid samples and corresponding bottom seawaters**

| **Hydrothermal vent** | **Guaymas Basin** | | **Mid-Cayman Von Damm**  **(Shallow)** | | **Mid-Cayman Piccard**  **(Deep)** | |
| --- | --- | --- | --- | --- | --- | --- |
| **Properties** | Theme Park | Bottom Seawater | East Summit | Bottom Seawater | Beebe 3 | Bottom Seawater |
| Temp (°C) | 315 | ~2.5 | 226 | 5 | 397 | 5 |
| Mn^2+^ | 0.24 | 0 | 6.50E-14 | 0 | 9.76E-07 | 0 |
| Fe^2+^ | 0.18 | 0 | 4.83E-13 | 0 | 2.07E-03 | 0 |
| Mg^2+^ | 0 | 53.5 | 0 | 52.4 | 0 | 52.4 |
| Cl^-^ | 637 | 538 | 658 | 545 | 358 | 545 |
| pH (at 25°C) | 5.9 | 8 | 5.56 | 8 | 3.17 | 8 |
| H_2_, aqueous | 3.4 | 0 | 19.2 | 0 | 20.7 | 0 |
| H_2_S | 5.98 | 0 | 3.2 | 0 | 12 | 0 |
| ∑CO_2_ | 61.1 | 2.18 | 2.78 | 2.2 | 26 | 2.2 |
| CO | 0 | 0 | 0 | 0 | 0 | 0 |
| CH_4_, aqueous | 63.4 | 0 | 2.84 | 0 | 0.123 | 0 |
| NH_4_^+^ | 13.6 | 0.0001 | 0.0179 | 0.0001 | 0.0347 | 0.0001 |
| O_2_, aqueous | 0 | 0.1^a^ | 0 | 0.25^b^ | 0 | 0.26^c^ |
| NO_3_^-^ | 0 | 0.037^a^ | 0 | 0.018^b^ | 0 | 0.018^c^ |
| NO_2_^-^ | 0 | 0.00003^a^ | 0 | -^b,d^ | 0 | -^c,d^ |
| N_2_^e^, aqueous | 0.48 | 0.58 | 0.48 | 0.58 | 0.48 | 0.58 |

All concentrations are in mmol/kg vent fluid or seawater. *In situ* pH values were calculated by using an equilibrium reaction path model that increased the temperature of the measured fluid to the original vent fluid temperature (*in situ* pH was transformed into the value at 25°C). The estimated temperature of bottom seawater is according to the previous reports^4^^,^ ^22^. The original chemical data is derived from the publications of Reeves et al.^3^ and Anantharaman et al^2^. For the end-member fluid concentrations of Lau Basin plumes, including Lau Basin Abe (A1 vent), Lau Basin Marine (MA1 vent), and Lau Basin Kilo Moana (KM1 vent), details are provided in Supplementary Table 2 in Anantharaman et al^2^.

a. Background concentrations at N 20°, W 105° of 1000m (NO_2_^-^) or 2000m (O_2_ and NO_3_^-^) depth in WOCE ATLAS (http://woceatlas.ucsd.edu/) VOLUME2 P18 section.

b. Background concentrations at N 18°, W 67° of 2000m depth in WOCE ATLAS VOLUME3 A22 section. NO_2_^-^ concentration is not applicable.

c. Background concentrations at N 18°, W 67° of 5000m depth in WOCE ATLAS VOLUME3 A22 section. NO_2_^-^ concentration is not applicable.

d. Treated as 0 in thermodynamic modeling.

e. Seawater dissolved N_2_^23^. Vent fluid dissolved N_2_ concentration is assumed to be 83% of seawater concentration^24^.

**Supplementary Table 3. Table of metabolic reactions and standard Gibbs free energies at 1, 25, and 100°C.** This table is adapted from the corresponding table from Anantharaman et al^4^.

| **Metabolism** | **Reaction** |  | **∆G°^a^ (kJ/mol)** | | |
| --- | --- | --- | --- | --- | --- |
|  |  | **e^-b^** | **1°C** | **25°C** | **100°C** |
| H_2_ oxidation (O_2_) | H_2_ + 0.5O_2_ → H_2_O | 2 | -265 | -264 | -260 |
| H_2_ oxidation  (NO_3_^-^ →N_2_) | NO_3_^-^ +2.5H_2_ + H^+^ → 0.5N_2_ + 3H_2_O | 5 | -637 | -637 | -635 |
| Methanotrophy | CH_4_ + 2O_2_ → HCO_3_^-^ + H^+^ + H_2_O | 8 | -828 | -825 | -810 |
| Sulfide oxidation  (O_2_)^c^ | HS^-^ + 2O_2_ → SO_4_^2-^ + H^+^ | 8 | -798 | -793 | -768 |
| Thiosulfate  oxidation (O_2_) | S_2_O_3_^2-^ + 2O_2_ + H_2_O → 2SO_4_^2-^ + 2H^+^ | 8 | -773 | -766 | -736 |
| Thiosulfate  oxidation (NO_3_^-^) | S_2_O_3_^2-^ + 1.6NO_3_^-^ + 0.2H_2_O → 2SO_4_^2-^ + 0.8N_2_ + 0.4H^+^ | 8 | -733 | -728 | -713 |
| Elemental Sulfur  oxidation (O_2_) | S^0^ + 1.5O_2_ + H_2_O → SO_4_^2-^ + 2H^+^ | 6 | -540 | -535 | -512 |
| H_2_ oxidation  (NO_3_^-^ →NO_2_^-^) | NO_3_^-^ +H_2_ → NO_2_^-^ + H_2_O | 2 | -177 | -177 | -175 |
| Elemental Sulfur  oxidation (NO_3_^-^) | S^0^ + 1.2NO_3_^-^ + 0.4H_2_O → SO_4_^2-^ + 0.6N_2_ + 0.8H^+^ | 6 | -510 | -507 | -494 |
| Sulfide oxidation  (NO_3_^-^) | NO_3_^-^ + HS^-^ + H^+^ → NO_2_^-^ + H_2_O + S^0^ | 2 | -170 | -170 | -172 |
| Iron reduction | H_2_ + 2Fe^3+^ → 2 Fe^2+^ + 2H^+^ | 2 | -89.6 | -92.5 | 100 |
| Ammonium  oxidation | NH_4_^+^ + 1.5O_2_ → 2H^+^ + NO_2_^−^ + H_2_O | 6 | -261 | -264 | -271 |
| H_2_ oxidation  (NH_4_^+^ →N_2_) | 0.5N_2_ + 1.5H_2_ + H^+^ → NH_4_^+^ | 3 | -217 | -216 | -207 |
| Sulfate reduction | SO_4_^2-^ + 4H_2_ + H^+^ → HS^-^ + 4H_2_O | 8 | -261 | -264 | -271 |
| Methanogenesis | 4H_2_ + HCO_3_^-^ H^+^ → CH_4_ +3H_2_O | 8 | -232 | -232 | -229 |
| Iron oxidation (O_2_)^c^ | Fe^2+^ + 0.25O_2_ + 2.5H_2_O → Fe(OH)_3,_s + 2H^+^ | 1 | -16.0 | -16.0 | -18.8 |
| Iron oxidation  (NO_3_^-^) | Fe^2+^ + 0.2NO_3_^-^ + 2.4H_2_O → Fe(OH)_3,_s + 0.1N_2_ + 1.8H^+^ | 1 | -11.0 | -11.3 | -15.8 |
| Manganese  oxidation | Mn^2+^ + 0.5O_2_ + H_2_O → MnO_2_,s + 2H^+^ | 2 | -5.65 | -5.71 | -4.62 |

a. Standard Gibbs free energies of reaction predicted by SUPCRT95^6^.
b. Number of electrons transferred during the reaction.
c. Reactions using particulate and aqueous phase e^-^ donors were predicted individually
(Sulfide and iron oxidation).
