## Supplementary Figures S1-S11 for "The sulfur cycle connects microbiomes and biogeochemistry in deep-sea hydrothermal plumes"

a

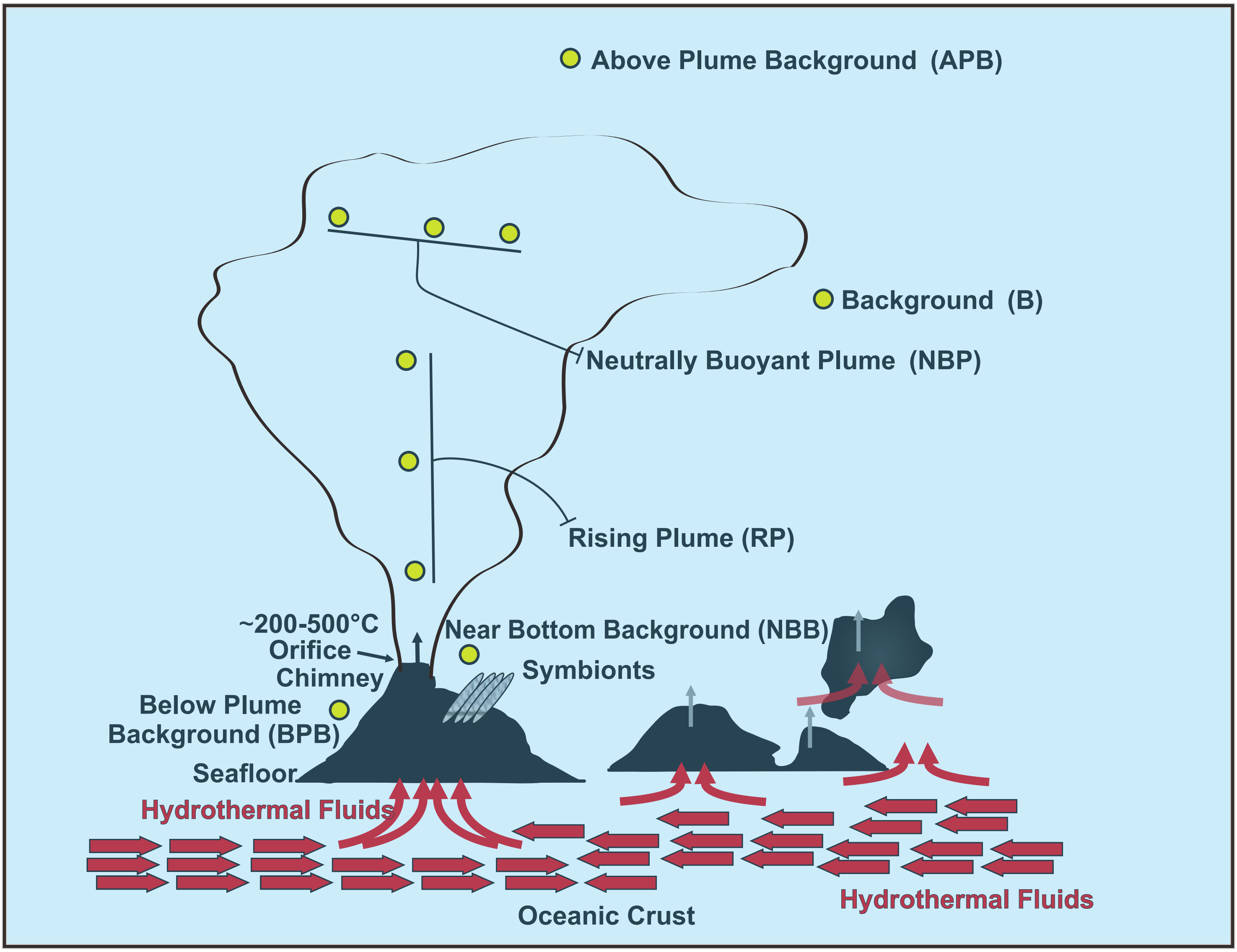


b

| | Sampling Places | Guaymas Basin | Mid-Cayman Von Damm (Shallow) | Mid-Cayman Piccard (Deep) | Lau Basin Kilo Moana | Lau Basin Tahi Moana | Lau Basin Abe | Lau Basin Tui Malia | Lau Basin Mariner |
|---|---|---|---|---|---|---|---|---|---|
| Background Samples(BS) | Above Plume Background (APB) | C | | | | D | | | |
| | Background (B) | | D C | C<br>D C | | | | | |
| | Near Bottom Background (NBB) | | C1 C2 | | | | D | | |
| | Below Plume Background (BPB) | | | | | | | | D |
| Plume Samples(PS) | Neutrally Buoyant Plume (NBP) | D C | | | D | | | | |
| | Rising Plume (RP) | | D1 C1 C2<br>D1 C1 C2 | C<br>D C<br>D C | D<br>D<br>D | D | D<br>D | D | D |

D→Metagenome C→Metatranscriptome C1 C2→Parallel samples D1 C1→Samples from the same filter, one for DNA, the other for cDNA

**Supplementary Figure S1. Schematic diagram of hydrothermal vent structure and sampling positions.** (**a**) Schematic diagram indicating detailed sample positions. (**b**) Summary table of DNA and cDNA sequencing libraries within this study.

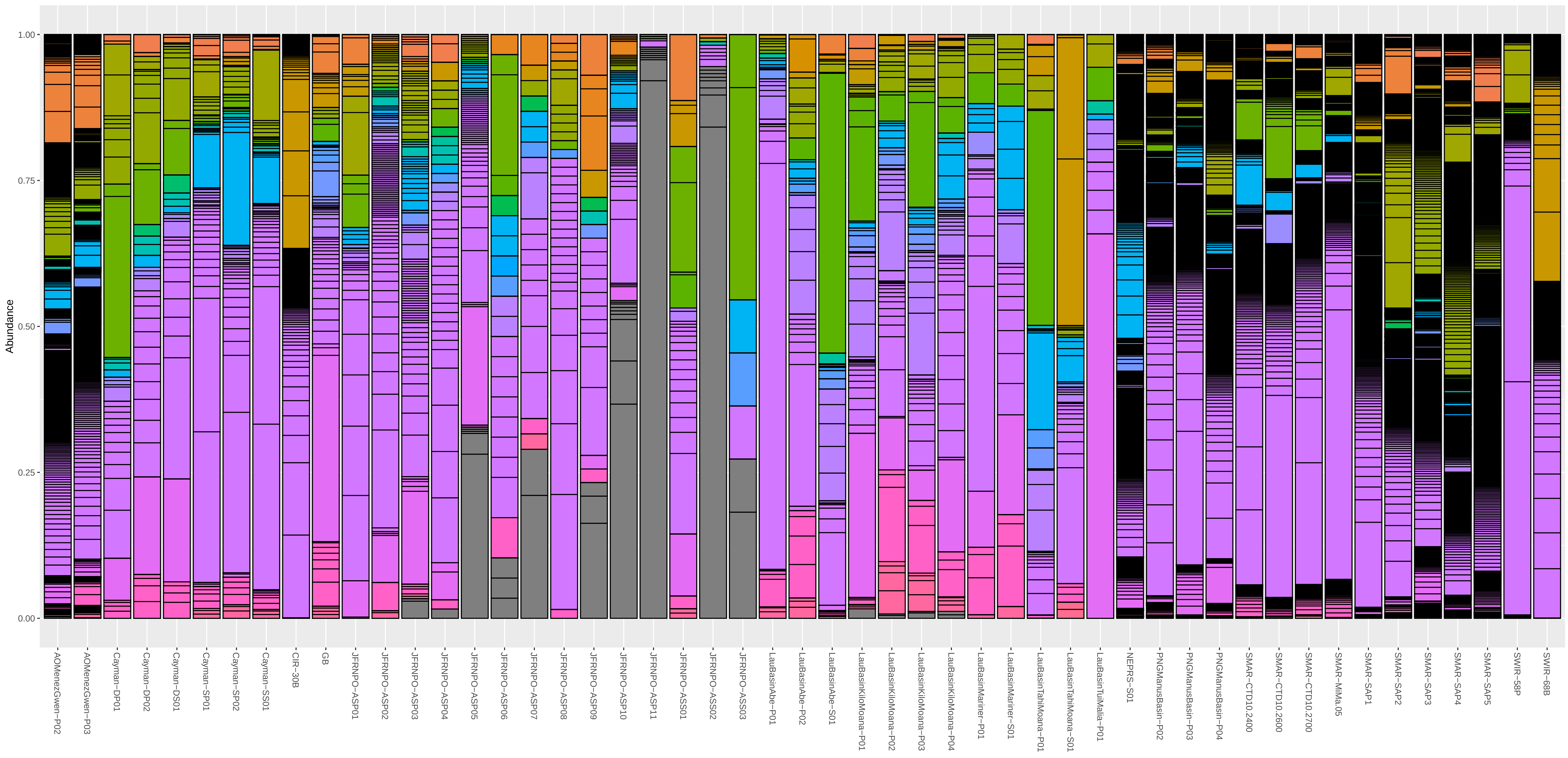


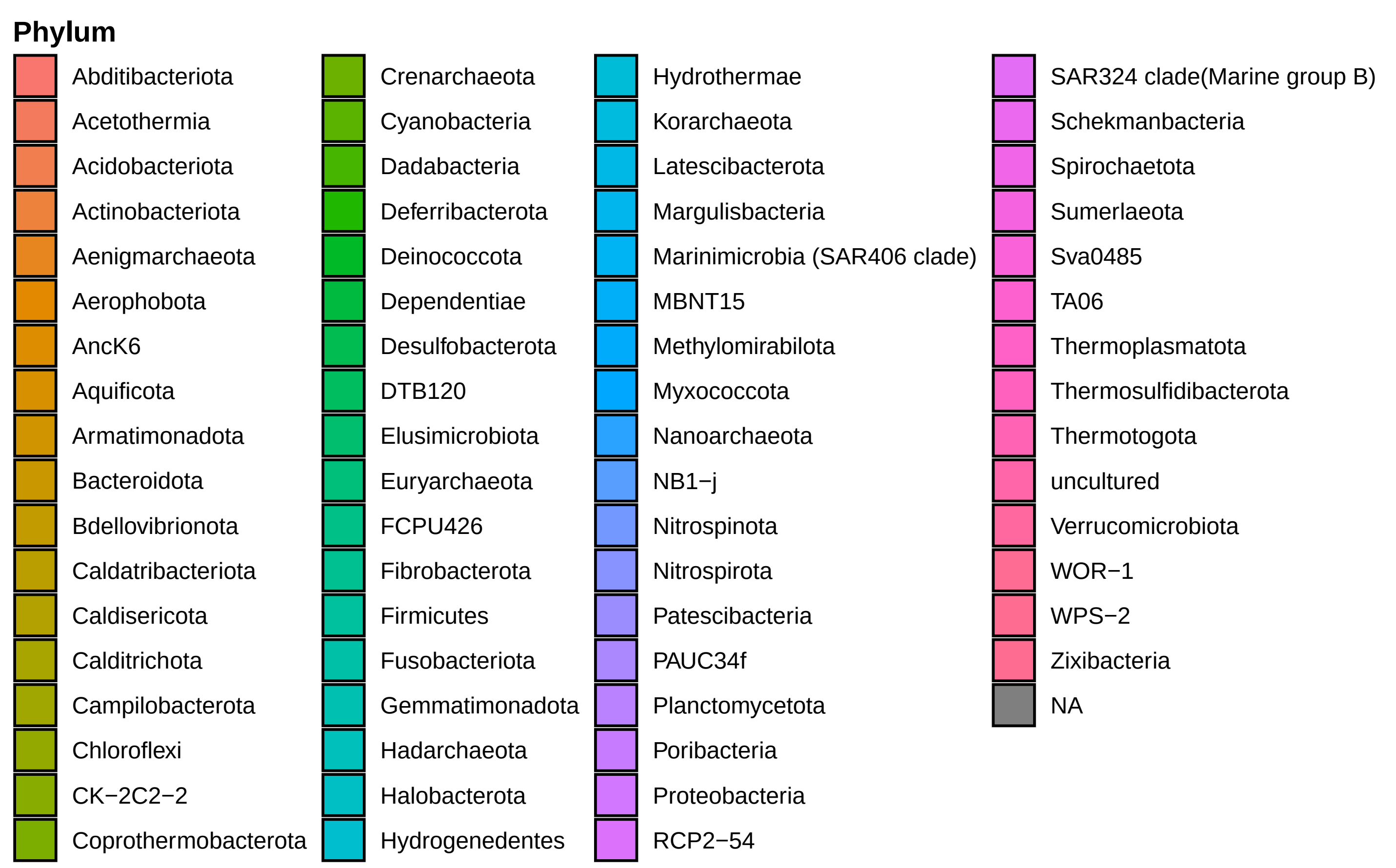


**Supplementary Figure S2. Compositional bar plot of hydrothermal plume and background microbial community at phylum level.** Microbial community analysis was based on 51 hydrothermal plume and background 16S rRNA gene datasets.

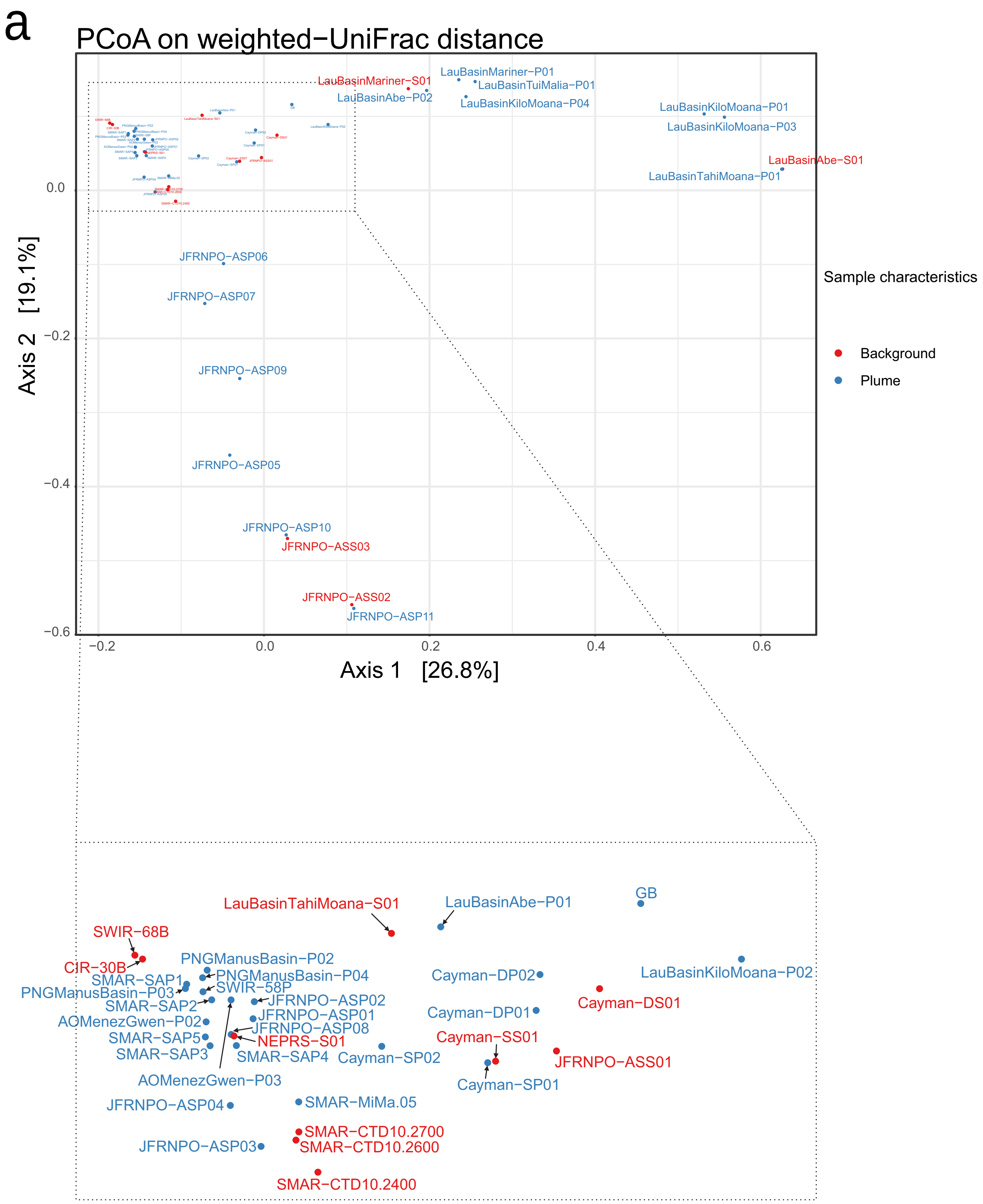
a
PCoA on weighted−UniFrac distance
Axis 1 [26.8%]
Axis 2 [19.1%]
Sample characteristics
Background
Plume
LauBasinMariner−S01
LauBasinMariner−P01
LauBasinTuiMalia−P01
LauBasinAbe−P02
LauBasinKiloMoana−P04
LauBasinKiloMoana−P01
LauBasinKiloMoana−P03
LauBasinAbe−S01
LauBasinTahiMoana−P01
JFRNPO−ASP06
JFRNPO−ASP07
JFRNPO−ASP09
JFRNPO−ASP05
JFRNPO−ASP10
JFRNPO−ASS03
JFRNPO−ASS02
JFRNPO−ASP11
LauBasinTahiMoana−S01
LauBasinAbe−P01
GB
SWIR−68B
CIR−30B
PNGManusBasin−P02
PNGManusBasin−P04
SMAR−SAP1
SWIR−58P
PNGManusBasin−P03
SMAR−SAP2
JFRNPO−ASP02
JFRNPO−ASP01
JFRNPO−ASP08
AOMenezGwen−P02
SMAR−SAP5
NEPRS−S01
SMAR−SAP3
SMAR−SAP4
AOMenezGwen−P03
Cayman−DP02
Cayman−DP01
Cayman−SS01
Cayman−SP02
Cayman−SP01
LauBasinKiloMoana−P02
Cayman−DS01
JFRNPO−ASS01
JFRNPO−ASP04
SMAR−MiMa.05
SMAR−CTD10.2700
SMAR−CTD10.2600
JFRNPO−ASP03
SMAR−CTD10.2400

b

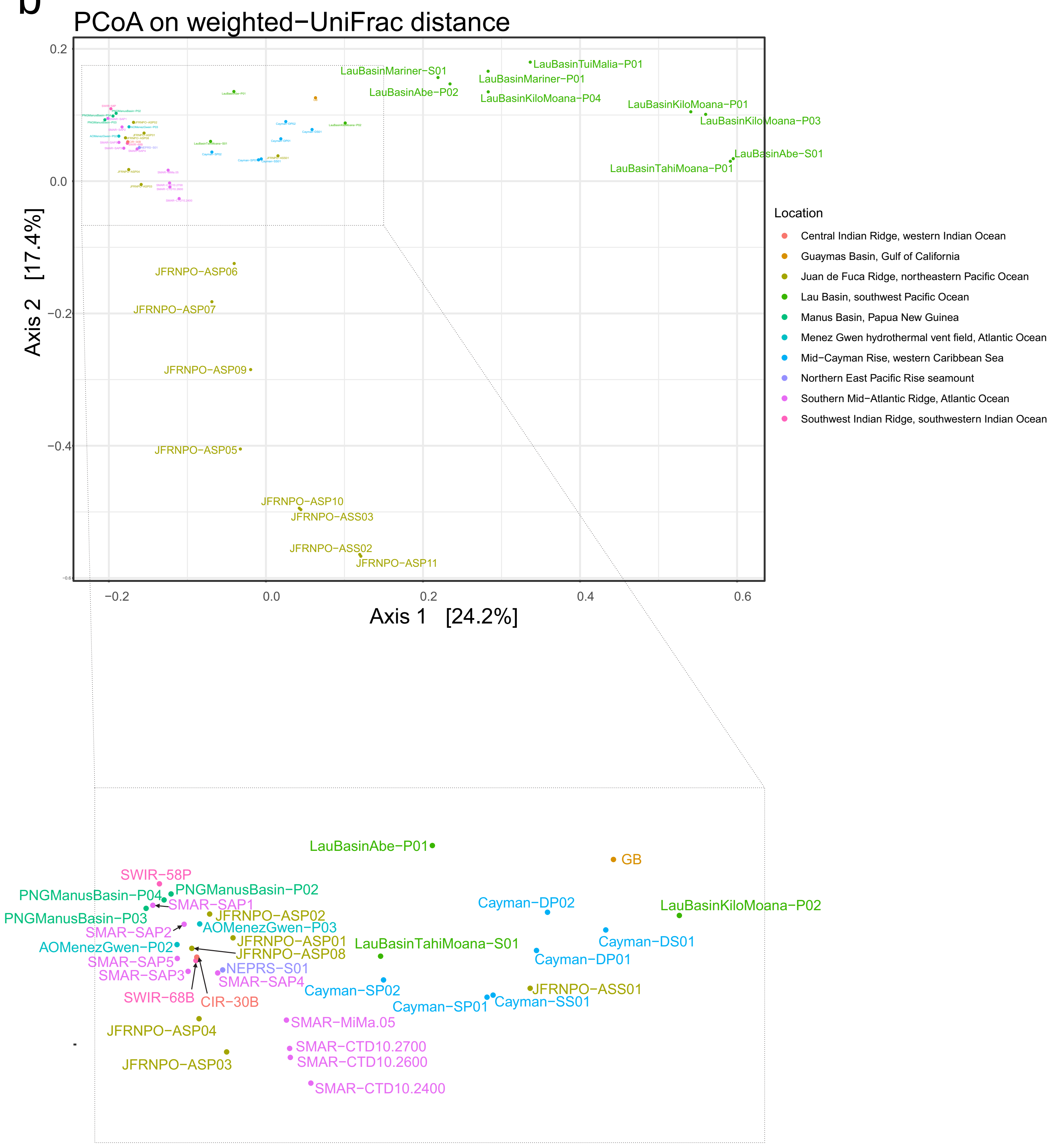
PCoA on weighted−UniFrac distance
Axis 1 [24.2%]
Axis 2 [17.4%]
0.2
0.0
−0.2
−0.4
−0.2
0.0
0.2
0.4
0.6
LauBasinMariner−S01
LauBasinTuiMalia−P01
LauBasinMariner−P01
LauBasinAbe−P02
LauBasinKiloMoana−P04
LauBasinKiloMoana−P01
LauBasinKiloMoana−P03
LauBasinAbe−S01
LauBasinTahiMoana−P01
JFRNPO−ASP06
JFRNPO−ASP07
JFRNPO−ASP09
JFRNPO−ASP05
JFRNPO−ASP10
JFRNPO−ASS03
JFRNPO−ASS02
JFRNPO−ASP11
Location
Central Indian Ridge, western Indian Ocean
Guaymas Basin, Gulf of California
Juan de Fuca Ridge, northeastern Pacific Ocean
Lau Basin, southwest Pacific Ocean
Manus Basin, Papua New Guinea
Menez Gwen hydrothermal vent field, Atlantic Ocean
Mid−Cayman Rise, western Caribbean Sea
Northern East Pacific Rise seamount
Southern Mid−Atlantic Ridge, Atlantic Ocean
Southwest Indian Ridge, southwestern Indian Ocean
LauBasinAbe−P01
GB
SWIR−58P
PNGManusBasin−P04
PNGManusBasin−P02
SMAR−SAP1
PNGManusBasin−P03
JFRNPO−ASP02
Cayman−DP02
LauBasinKiloMoana−P02
SMAR−SAP2
AOMenezGwen−P03
JFRNPO−ASP01
LauBasinTahiMoana−S01
Cayman−DS01
AOMenezGwen−P02
JFRNPO−ASP08
Cayman−DP01
SMAR−SAP5
NEPRS−S01
SMAR−SAP3
SMAR−SAP4
Cayman−SP02
JFRNPO−ASS01
SWIR−68B
CIR−30B
Cayman−SP01
Cayman−SS01
SMAR−MiMa.05
JFRNPO−ASP04
SMAR−CTD10.2700
JFRNPO−ASP03
SMAR−CTD10.2600
SMAR−CTD10.2400

C

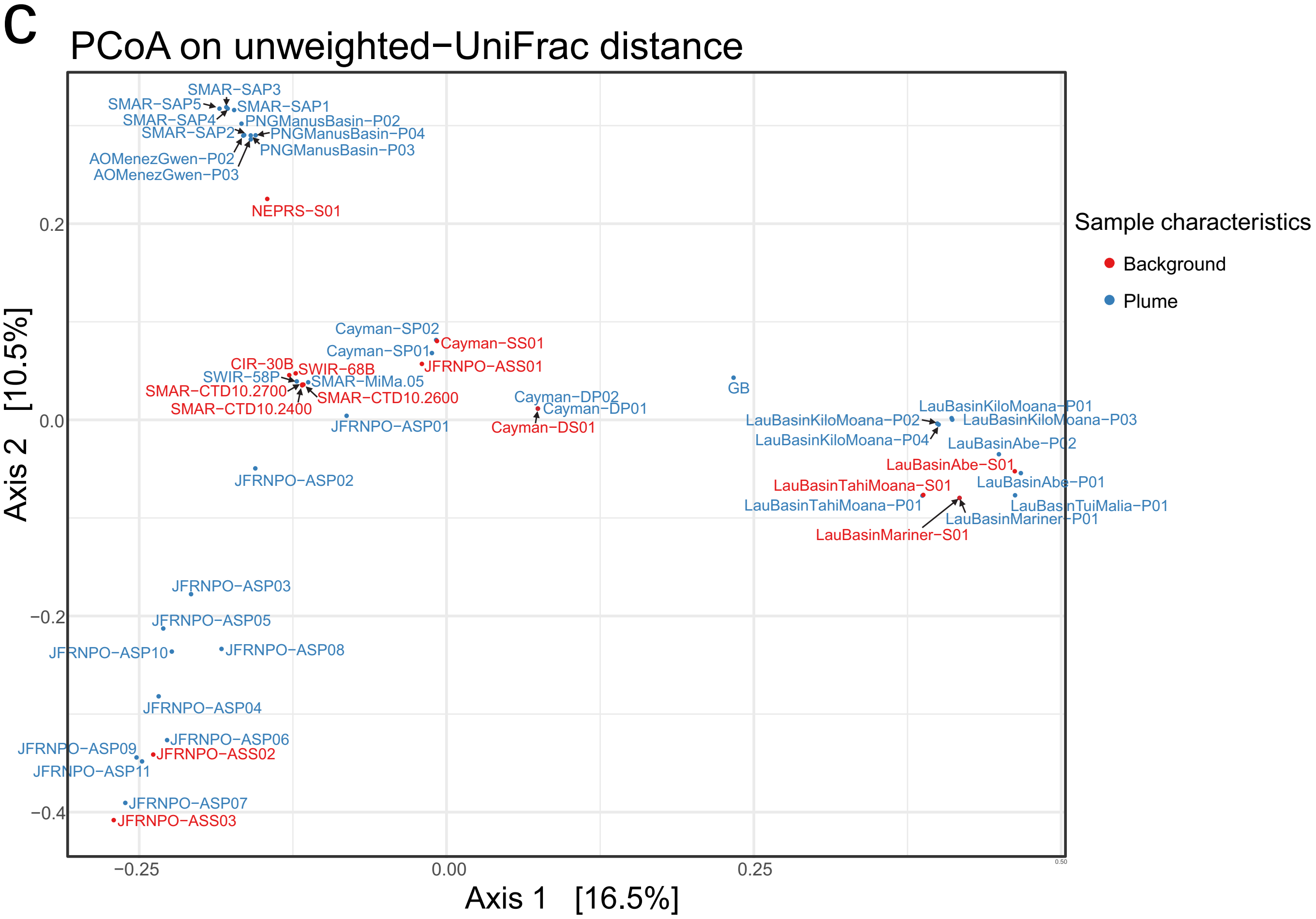

PCoA on unweighted−UniFrac distance
Axis 1 [16.5%]
Axis 2 [10.5%]
Sample characteristics
Background
Plume
SMAR−SAP3
SMAR−SAP5
SMAR−SAP1
SMAR−SAP4
PNGManusBasin−P02
SMAR−SAP2
PNGManusBasin−P04
PNGManusBasin−P03
AOMenezGwen−P02
AOMenezGwen−P03
NEPRS−S01
Cayman−SP02
Cayman−SS01
Cayman−SP01
JFRNPO−ASS01
CIR−30B
SWIR−68B
SWIR−58P
SMAR−MiMa.05
SMAR−CTD10.2700
SMAR−CTD10.2600
SMAR−CTD10.2400
JFRNPO−ASP01
Cayman−DP02
Cayman−DP01
Cayman−DS01
GB
LauBasinKiloMoana−P01
LauBasinKiloMoana−P02
LauBasinKiloMoana−P03
LauBasinKiloMoana−P04
LauBasinAbe−P02
LauBasinAbe−S01
LauBasinTahiMoana−S01
LauBasinAbe−P01
LauBasinTahiMoana−P01
LauBasinTuiMalia−P01
LauBasinMariner−P01
LauBasinMariner−S01
JFRNPO−ASP02
JFRNPO−ASP03
JFRNPO−ASP05
JFRNPO−ASP10
JFRNPO−ASP08
JFRNPO−ASP04
JFRNPO−ASP06
JFRNPO−ASP09
JFRNPO−ASS02
JFRNPO−ASP11
JFRNPO−ASP07
JFRNPO−ASS03
0.2
0.0
−0.2
−0.4
−0.25
0.00
0.25

d

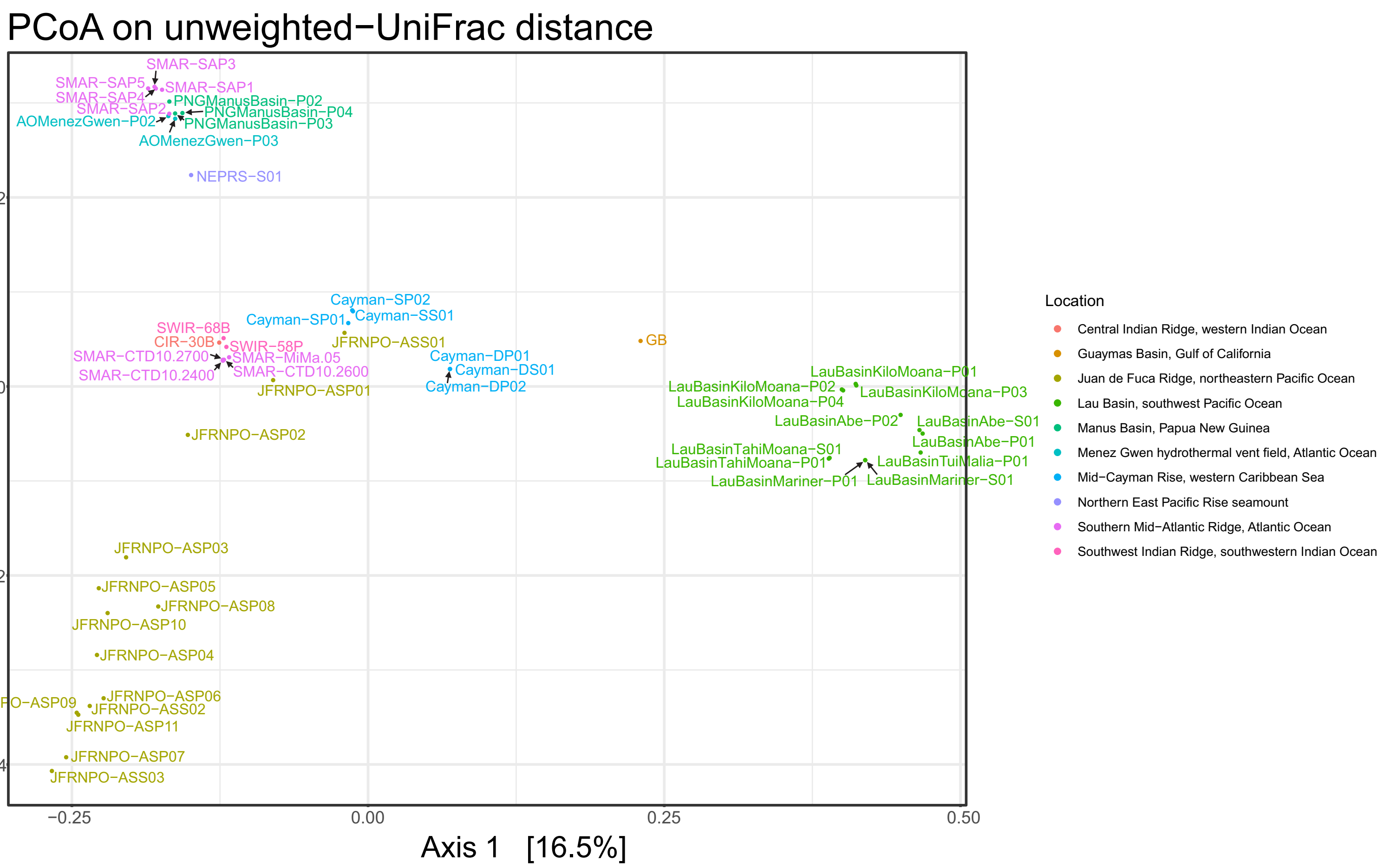


**Supplementary Figure S3. PCoA diagrams of global hydrothermal plume and background microbial community based on 16S rRNA gene.** Four subpanels contained PCoA diagrams each based on weighted/unweighted Unifrac distance (a, b based on weighted Unifrac distance; c, d based on unweighted Unifrac distance) and labeled by location/sample characteristics (plume or background) (a, c labeled by sample characteristics; b, d labeled by location characteristics).

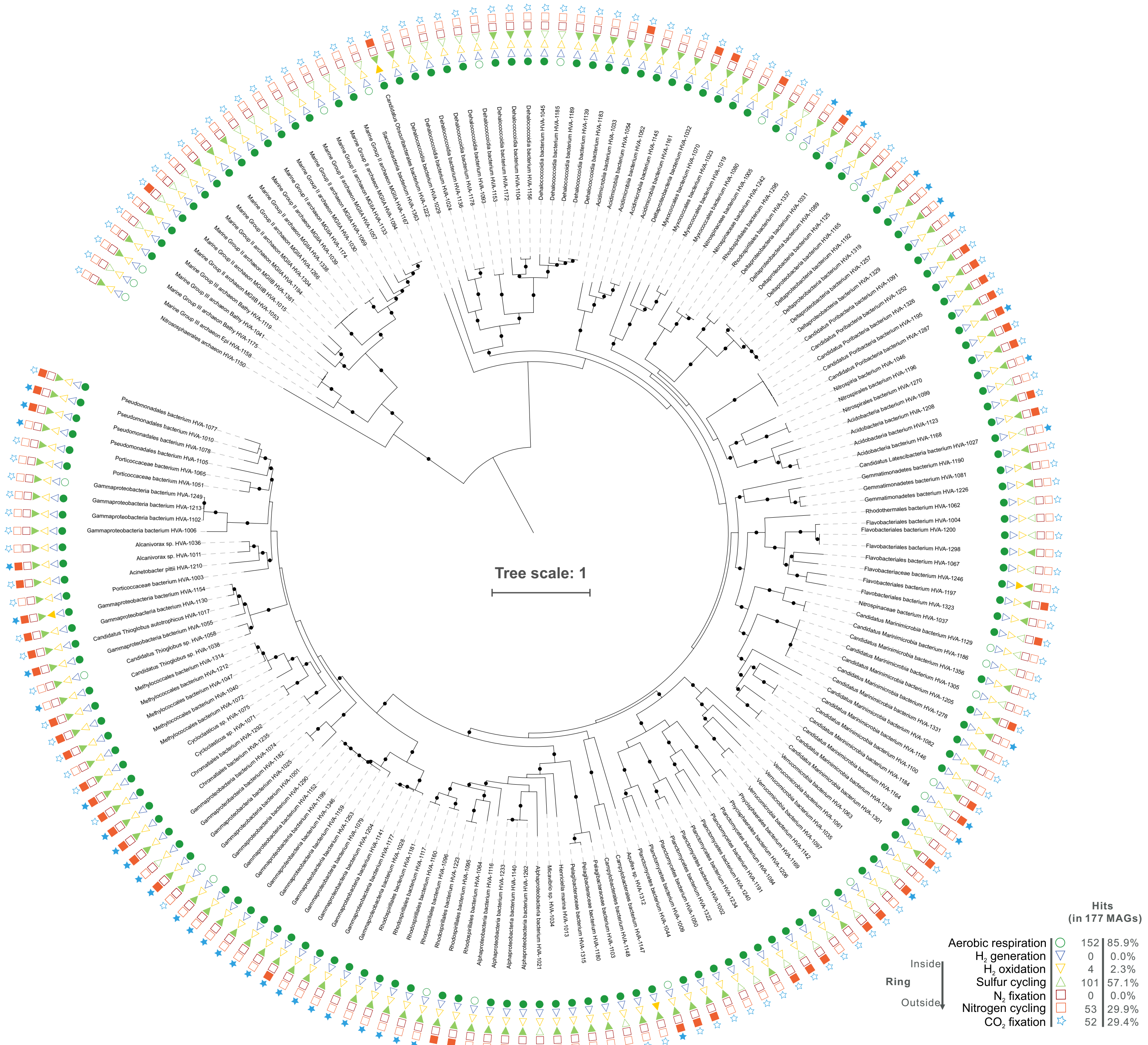


**Supplementary Figure S4. Phylogenetic tree of MAGs based on concatenated 16 ribosomal proteins.** Only the nodes with ultrafast bootstrap (UFBoot) support values over 90% were labeled with black dots. The functional traits for each MAG were parsed and labeled in the tree according to the HMM scan result. Filled shapes indicate presence of functional traits, blank shapes indicate absence of functional traits.

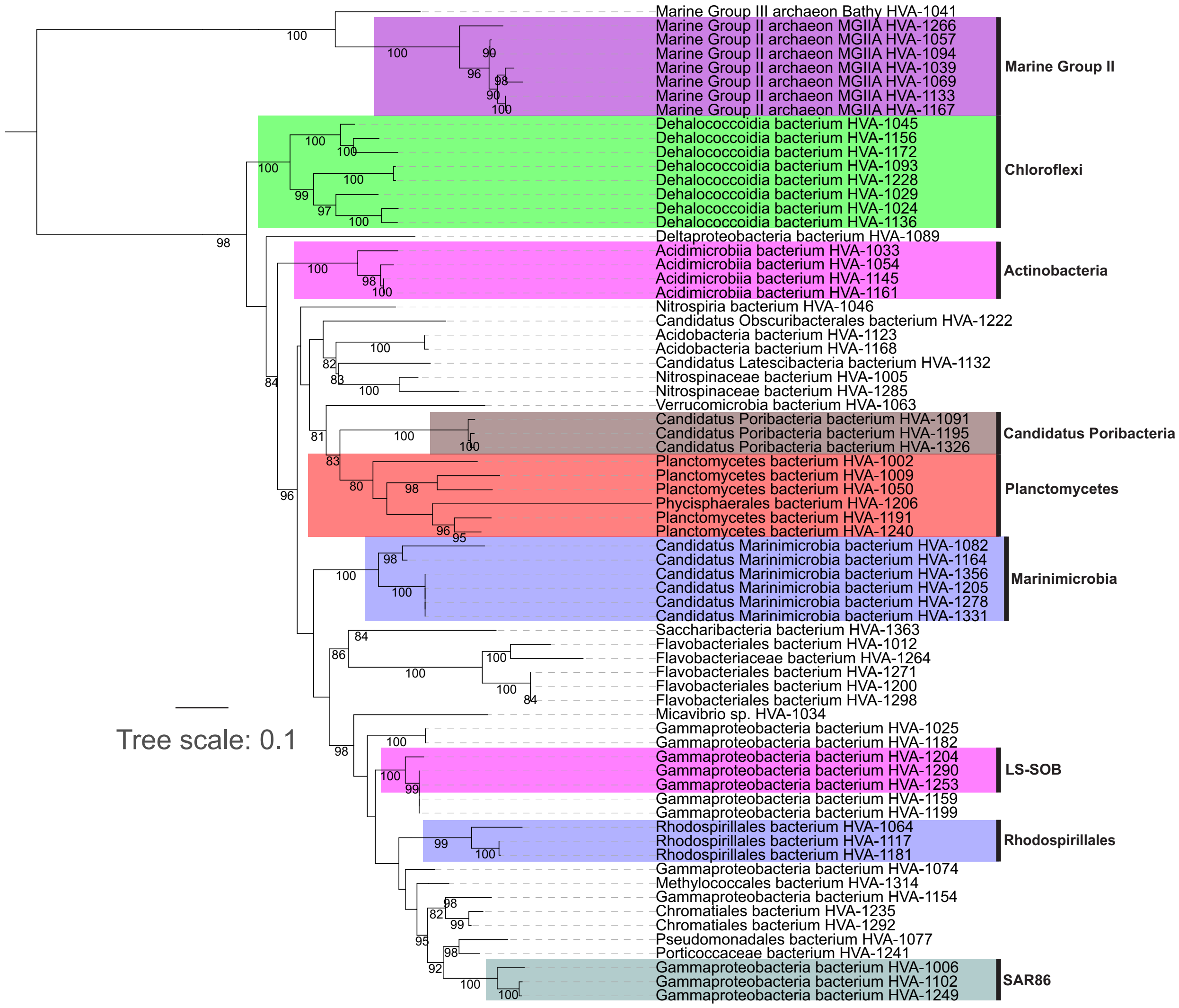


**Supplementary Figure S5. Phylogenetic tree based on 16S rRNA gene from each MAG.** The bootstrap (UFBoot) support values were labeled to each node (only showing those > 80%). Taxonomic labels were according to both SILVA_128_SSUParc_tax_silva database BLAST result and MAG 16RP phylogenetic tree.

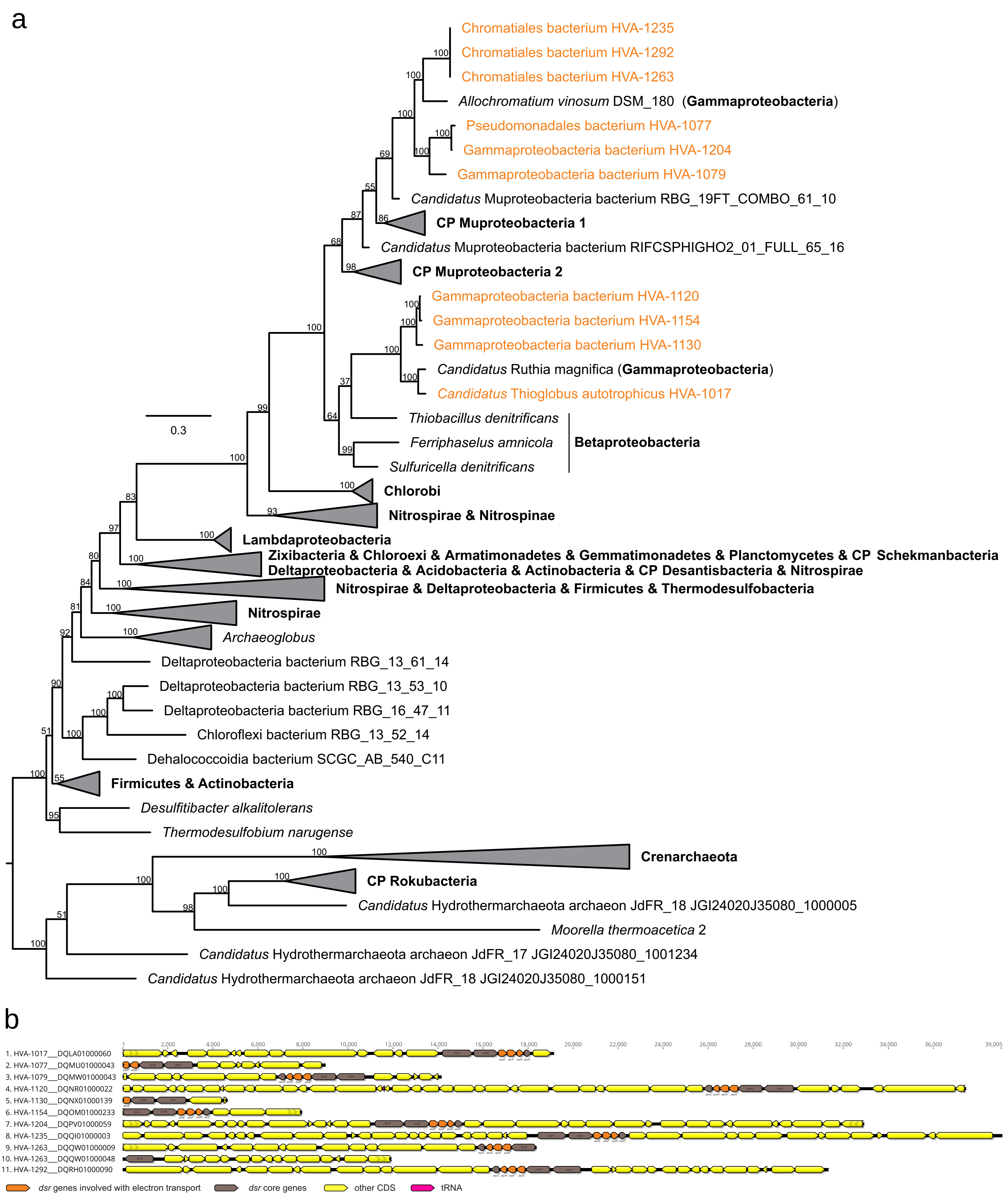


**Supplementary Figure S6. Phylogenetic tree of concatenated *dsrAB* encoding proteins and gene structure figure of *dsr* containing scaffolds.** (**a**) Phylogenetic tree of concatenated *dsrAB* encoding proteins. DsrA and DsrB were aligned with reference sequences independently and concatenated. The concatenated protein alignment was trimmed with gap threshold of 25% using trimAl v1.2. The phylogenetic tree was reconstructed by IQ-TREE v1.6.9 with settings as described in the methods. UFBoot bootstrap values were labeled at each node. Genomes from this study were highlighted in yellow. (**b**) Gene structure figure of *dsr* containing scaffolds. The assignment of each *dsr* gene component was confirmed by combining BLAST result in NCBI database and position in the *dsr* gene operon. This figure was visualized in Geneious Prime v2020.2.3.

a

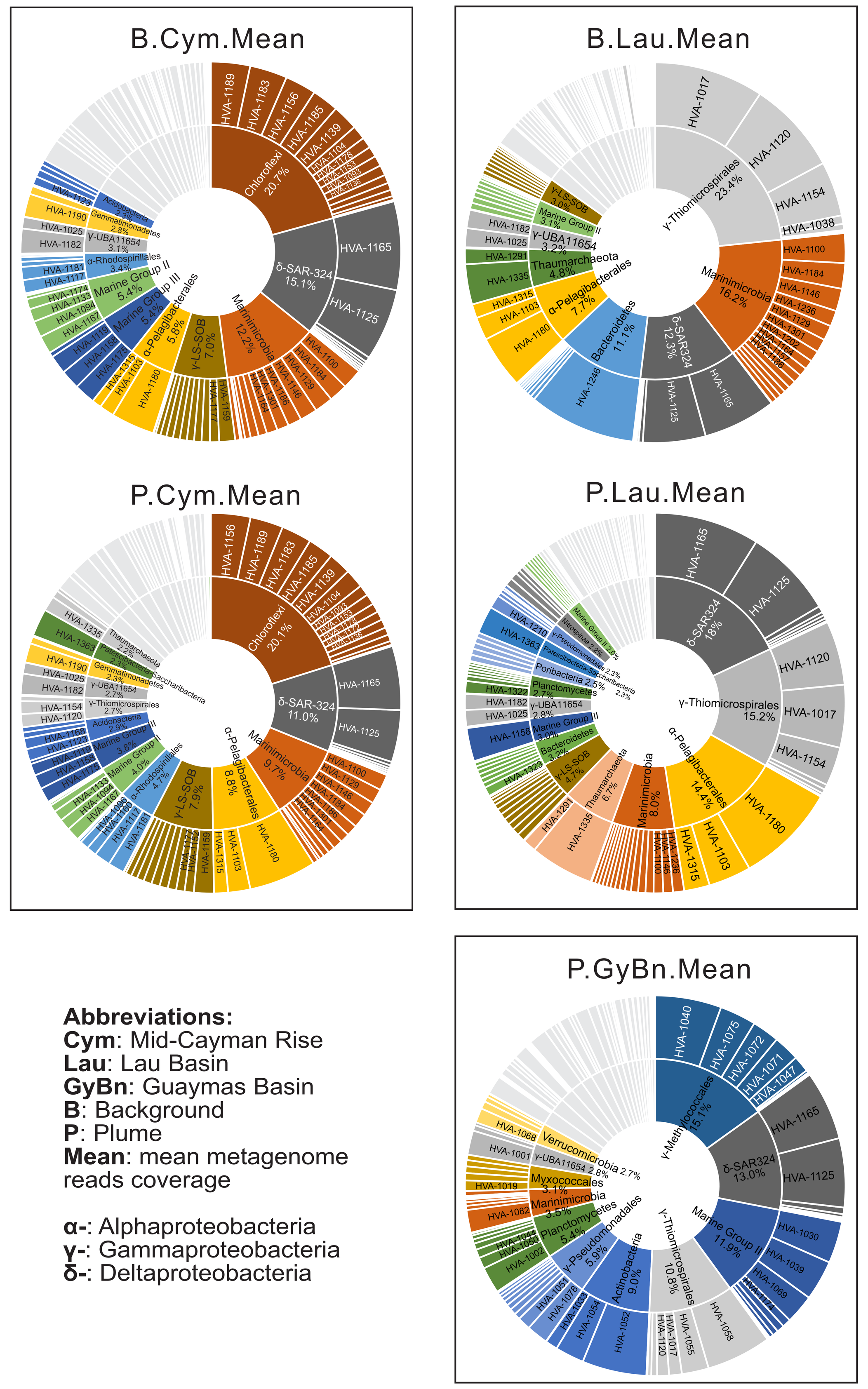


| Microbial Group | Log2 Fold Change | adjusted *P*-value | MAG MetaG Cov. Mean value | |
|---|---|---|---|---|
| **B.Cym-v-B.Lau** | | | B.Cym.mean | B.Lau.mean |
| γ-Thiomicrospirales | 4.53 | 2.54E-04 | 1.0% | 23.4% |
| Bacteroidetes | 4.48 | 4.55E-03 | 0.5% | 11.1% |
| Chloroflexi | -3.85 | 4.17E-03 | 20.7% | 1.5% |
| **P.Cym-v-P.Lau** | | | P.Cym.mean | P.Lau.mean |
| γ-Thiomicrospirales | 2.70 | 5.39E-06 | 2.7% | 15.2% |
| Thaumarchaeota | 2.01 | 7.03E-06 | 2.2% | 6.7% |
| Chloroflexi | -3.48 | 1.88E-06 | 20.1% | 1.5% |
| **P.Cym-v-P.GyBn** | | | P.Cym.mean | P.GyBn.mean |
| γ-Methylococcales | 8.09 | 1.80E-88 | 0.1% | 15.1% |
| γ-Pseudomonadales | 3.27 | 6.19E-05 | 0.7% | 5.9% |
| Planctomycetes | 2.73 | 3.62E-02 | 0.8% | 5.4% |
| γ-Thiomicrospirales | 2.09 | 4.85E-06 | 2.7% | 10.8% |
| Marine Group II | 1.65 | 5.37E-03 | 4.0% | 11.9% |
| Marinimicrobia | -1.38 | 3.49E-02 | 9.7% | 3.5% |
| γ-LS-SOB | -2.35 | 3.08E-18 | 7.9% | 1.4% |
| α-Pelagibacterales | -3.55 | 2.16E-24 | 8.8% | 0.7% |
| Chloroflexi | -3.70 | 8.08E-09 | 20.1% | 1.5% |
| **P.GyBn-v-P.Lau** | | | P.GyBn.mean | P.Lau.mean |
| α-Pelagibacterales | 4.63 | 8.64E-04 | 0.7% | 14.4% |
| Thaumarchaeota | 3.37 | 1.56E-04 | 0.8% | 6.7% |
| γ-Methylococcales | -3.88 | 3.22E-03 | 15.1% | 0.9% |
| Actinobacteria | -7.82 | 2.06E-155 | 9.0% | 0.0% |

b

| Function | Gene name/ Subfunction/dbCAN2/ MEROPS | Target ID | Log2 Fold Change | adjusted *P*-value | Function trait MetaG Cov. Mean value | |
|---|---|---|---|---|---|---|
| | | **B.Cym-v-B.Lau** | | | B.Cym.mean | B.Lau.mean |
| Arsenate reduction | arsC | TIGR00014 | 1.35 | 8.48E-03 | 0.2% | 0.6% |
| | | **P.Cym-v-P.Lau** | | | P.Cym.mean | P.Lau.mean |
| Arsenate reduction | arsC | TIGR00014 | 1.09 | 2.02E-07 | 0.3% | 0.6% |
| long-chain acyl-CoA synthetase ($C_6$+) | ACSL | KEGG.B20 | 0.59 | 9.67E-09 | 1.2% | 1.6% |
| Halogenated compounds breakdown | Haloacid dehydrogenase | TIGR01428 | -1.08 | 2.06E-02 | 1.7% | 0.7% |
| CO oxidation | coxS | carbon_monoxide_dehydrogenase_coxS | -1.51 | 3.21E-03 | 2.1% | 0.7% |
| CO oxidation | coxL | TIGR02416 | -1.94 | 2.20E-03 | 2.9% | 0.7% |
| Methanol oxidation | pqq EC 1.1.2.7 | methanol_dehydrogenase_pqq_xoxF_mxaF | -2.06 | 2.84E-02 | 0.6% | 0.1% |
| | | **P.Cym-v-P.GyBn** | | | P.Cym.mean | P.GyBn.mean |
| Benzene degradation, benzene => catechol | dmpKLMNOP | KEGG.B05 | 9.03 | 3.21E-16 | 0.0% | 1.2% |
| Toluene degradation, toluene => benzoate | tmoABCDEF | KEGG.B01 | 3.59 | 9.63E-22 | 0.1% | 0.8% |
| Iron oxidation | cyc1 | cyc1 | 0.97 | 2.75E-03 | 0.3% | 0.5% |
| Sulfate reduction | cysC | TIGR00455 | 0.74 | 1.83E-02 | 0.3% | 0.6% |
| CO oxidation | coxL | TIGR02416 | -1.85 | 4.73E-02 | 2.9% | 0.8% |
| Halogenated compounds breakdown | Haloacid dehydrogenase | TIGR01428 | -2.88 | 2.59E-07 | 1.7% | 0.2% |
| | | **P.GyBn-v-P.Lau** | | | P.GyBn.mean | P.Lau.mean |
| long-chain acyl-CoA synthetase ($C_6$+) | ACSL | KEGG.B20 | 0.76 | 2.30E-03 | 1.2% | 1.6% |
| Toluene degradation, toluene => benzoate | tmoABCDEF | KEGG.B01 | -8.61 | 4.55E-32 | 0.8% | 0.0% |

**Supplementary Figure S7. Sunburst figures and tables representing the comparison of MAG taxonomic composition and abundance and functional trait of samples from three hydrothermal environments based on metagenomic read mapping results.** (**a**) sunburst figures and table for comparison of MAG taxonomic composition. The mean MetaG MAG coverage values of background (B) and plume (P) were used to draw the sunburst figure. DESeq-based statistical analysis on the abundance difference of MAGs that are affiliated to certain microbial groups indicated significantly differentiated microbial groups between the comparisons of each two out of three environments. Only the microbial groups with > 5% mean coverage percentage in either one of the environments were listed with the corresponding Log2 Fold Change values and adjusted *P*-values (by nbinomWaldTest). (**b**) Table for comparison of functional traits. DESeq-based statistical analysis on the abundance of functional traits indicated significantly differentiated functional traits between the comparisons of each two out of three environments. The corresponding Log2Fold Change values and adjusted *P*-values (by nbinomWaldTest) were labeled for each identified functional trait.

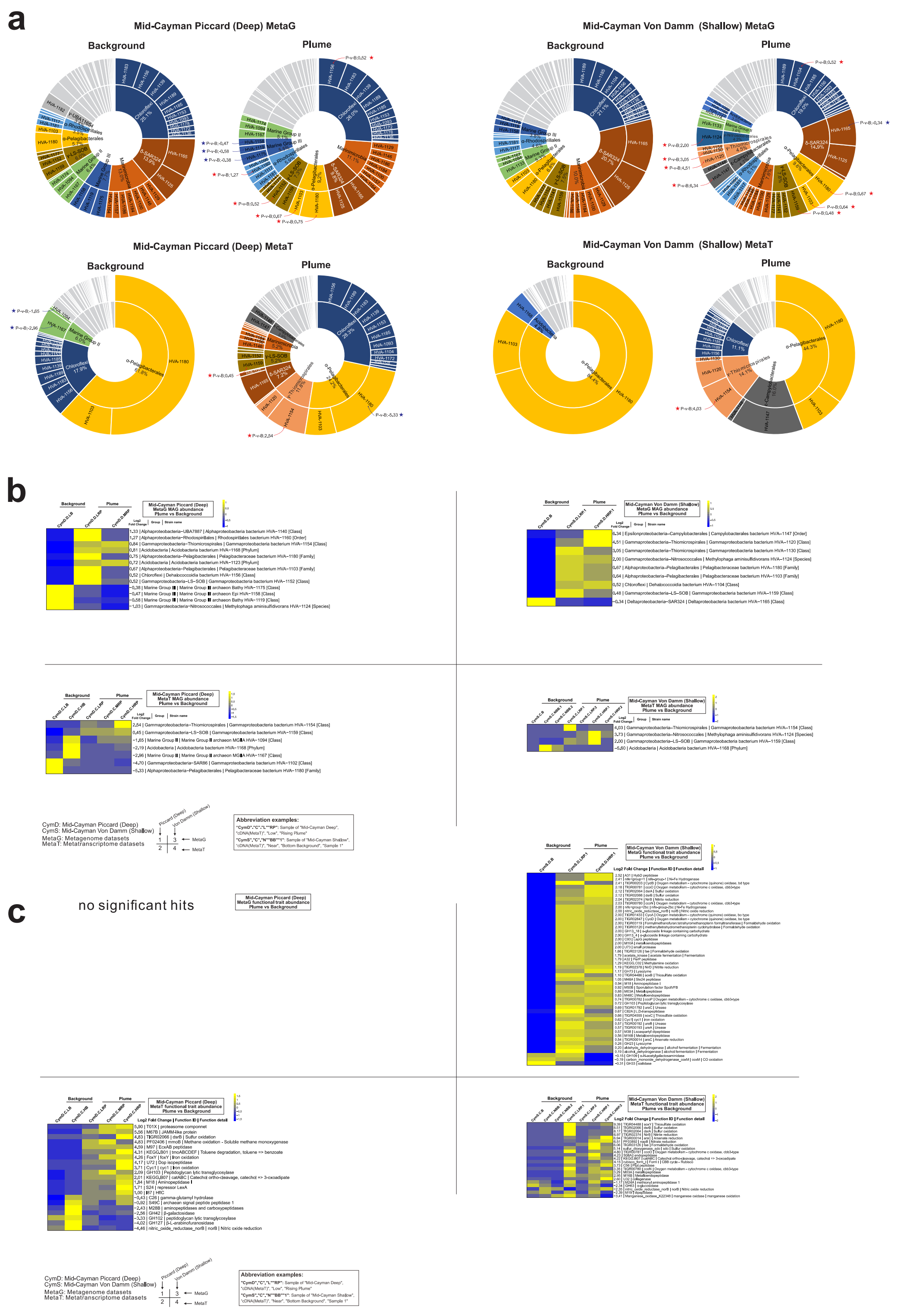


**Supplementary Figure S8. Sunburst diagrams and heatmaps representing DESeq result of MAG and functional traits between plume and background samples based on metagenome and metatranscriptome datasets from Mid-Cayman hydrothermal environments.** (**a**) Sunburst diagrams of MAGs based on metagenome and metatranscriptome. DESeq-based statistical analysis on the abundance and active abundance difference of MAGs from P-v-B (Plume vs Background) comparisons indicated MAGs with significant adjusted *P*-values ($P < 0.05$, red or blue star labeled) have differentiated abundances/active abundances in different hydrothermal eco-niches. Red stars indicated positive Log2 Fold Change, while blue stars indicated negative Log2 Fold Change; Log2 Fold Change values were also labeled with the stars accordingly. Only microbial groups with > 3% relative abundance in sunburst diagrams were labeled, while minor microbial groups were grey-colored. (**b**) Heatmap indicating MAG abundance difference based on metagenome and metatranscriptome. (**c**) Heatmap indicating functional trait abundance difference based on metagenome and metatranscriptome. Corresponding abundances shown here were of significant adjusted *P*-values (by nbinomWaldTest). Log2 Fold Change values were labeled accordingly. Relative abundance at row was normalized by removing the mean (centering) and dividing by the standard deviation (scaling). Resulted Row Z-score bars were presented accordingly in individual subpanels.

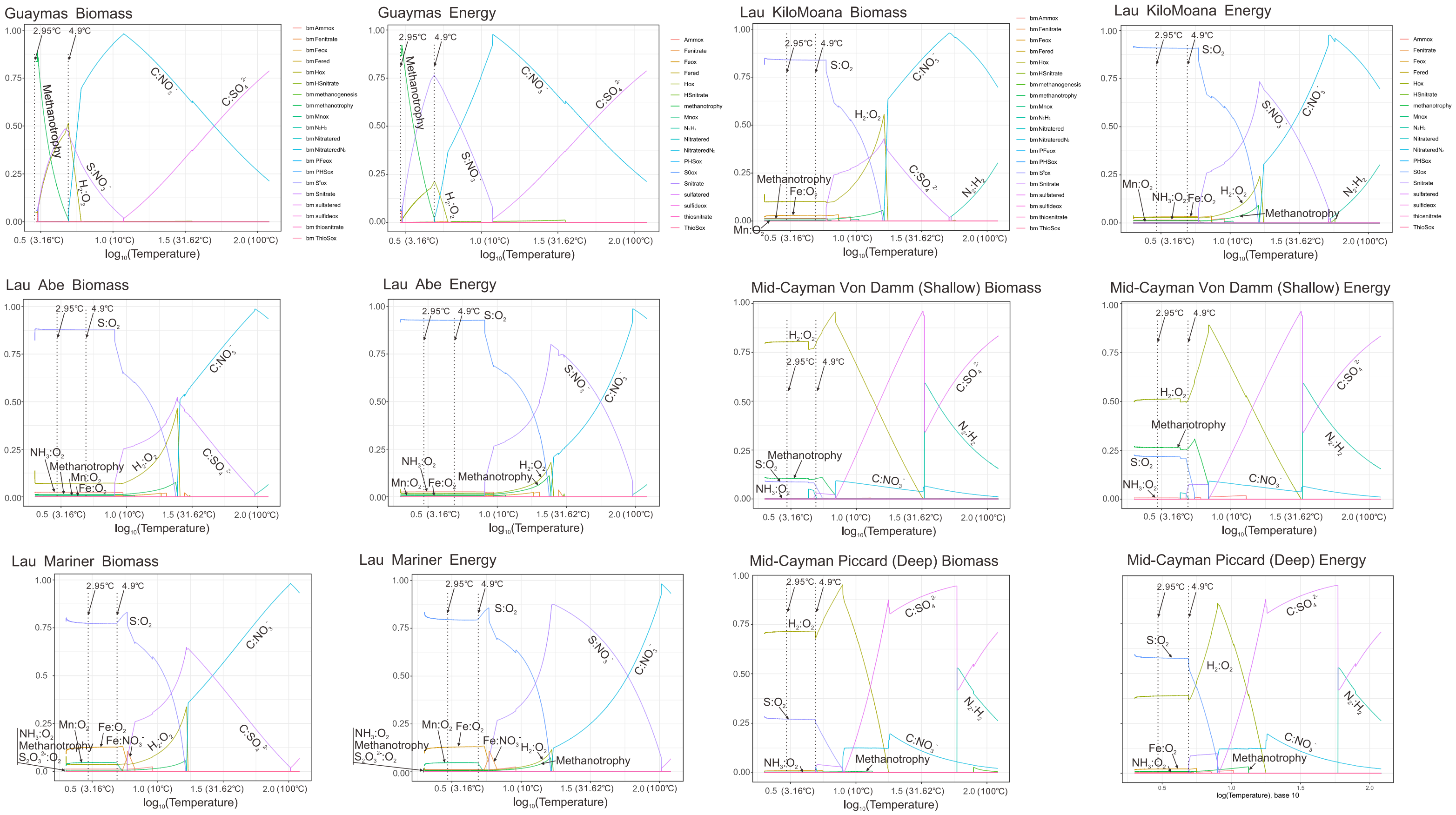


**Supplementary Figure S9. Thermodynamic estimation of available free energies and biomasses from reactions of electron donors in various hydrothermal plumes.** The estimated biomasses and free energies of individual environments were normalized to percentage fractions. Dotted lines (one at 2.95℃ and one at 4.9℃) showed two temperatures that we picked to conduct the biomass and free energy estimations for representing upper and lower plume temperatures. The abbreviation of reaction was labelled as: "$S:O_2$" standing for the reaction of sulfur as the electron donor and oxygen as the electron acceptor.

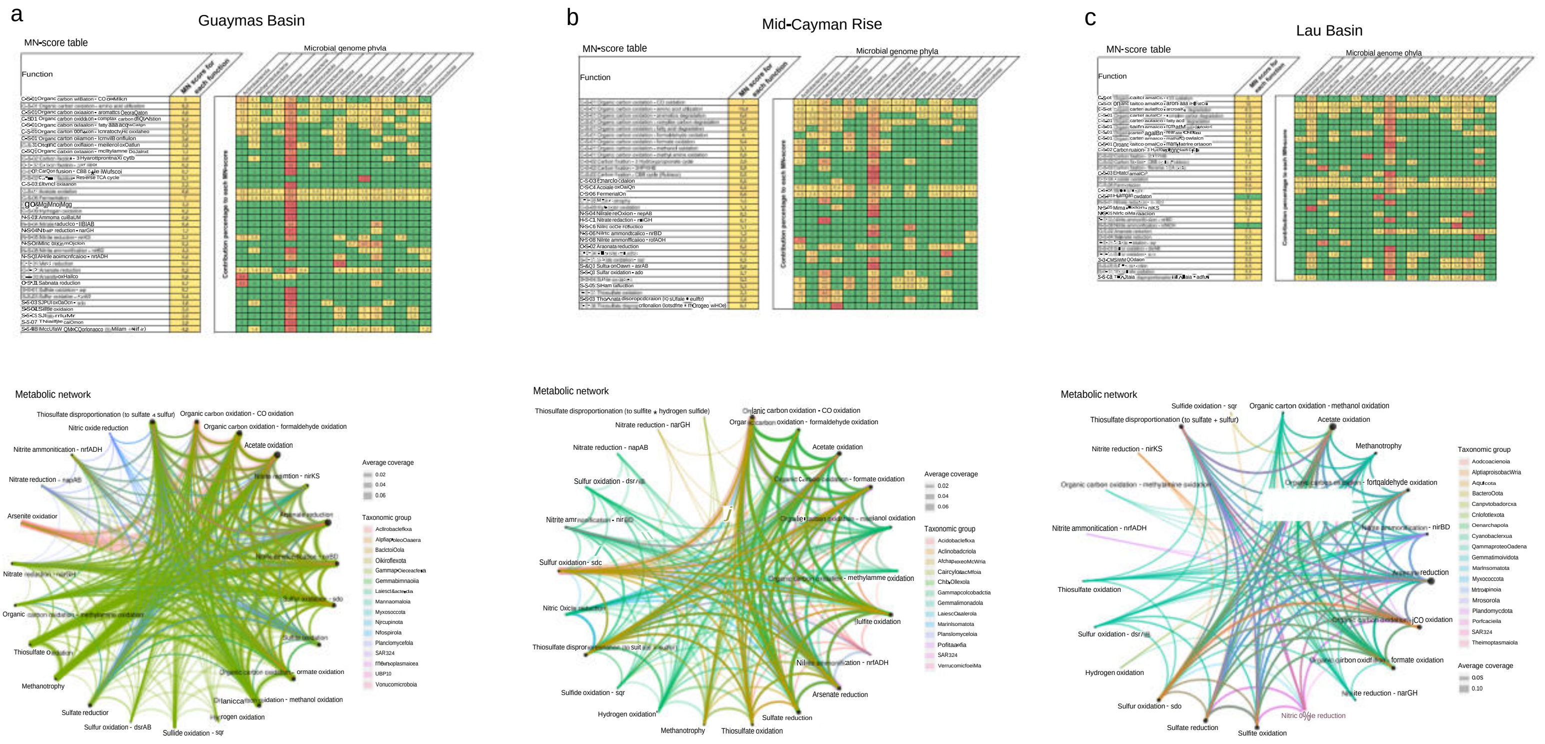


**Supplementary Figure S10. MW-scores (metabolic weight scores) and functional network diagrams of three hydrothermal vent sites. (a)** MW-score table and functional network diagram of Guaymas Basin. **(b)** MW-score table and functional network diagram of Mid-Cayman Rise. (**c**) MW-score table and functional network diagram of Lau Basin. MW-score tables and functional networking diagrams were generated by METABOLIC-C software using plume metagenomic reads as inputs. MW-score reflects abundance fraction of individual metabolic steps within the whole community. In each MW-score table, MW-scores are given in a separated column, and the rest part of the table indicates the contribution percentage to each MW-score of the genomes within the community as grouped by each phylum. From METABOLIC-C standard MW-score results, a group of metabolic cycling steps that are important on reflecting the plume substrate metabolisms were selected to make functional network diagrams. In each diagram, the size of a node is proportional to the gene coverage associated with the metabolic/biogeochemical cycling step. The thickness of the edge was depicted according to the average of gene coverage values of the two connected biogeochemical cycling steps. The color of the edges was assigned based on the taxonomy of the represented genome.

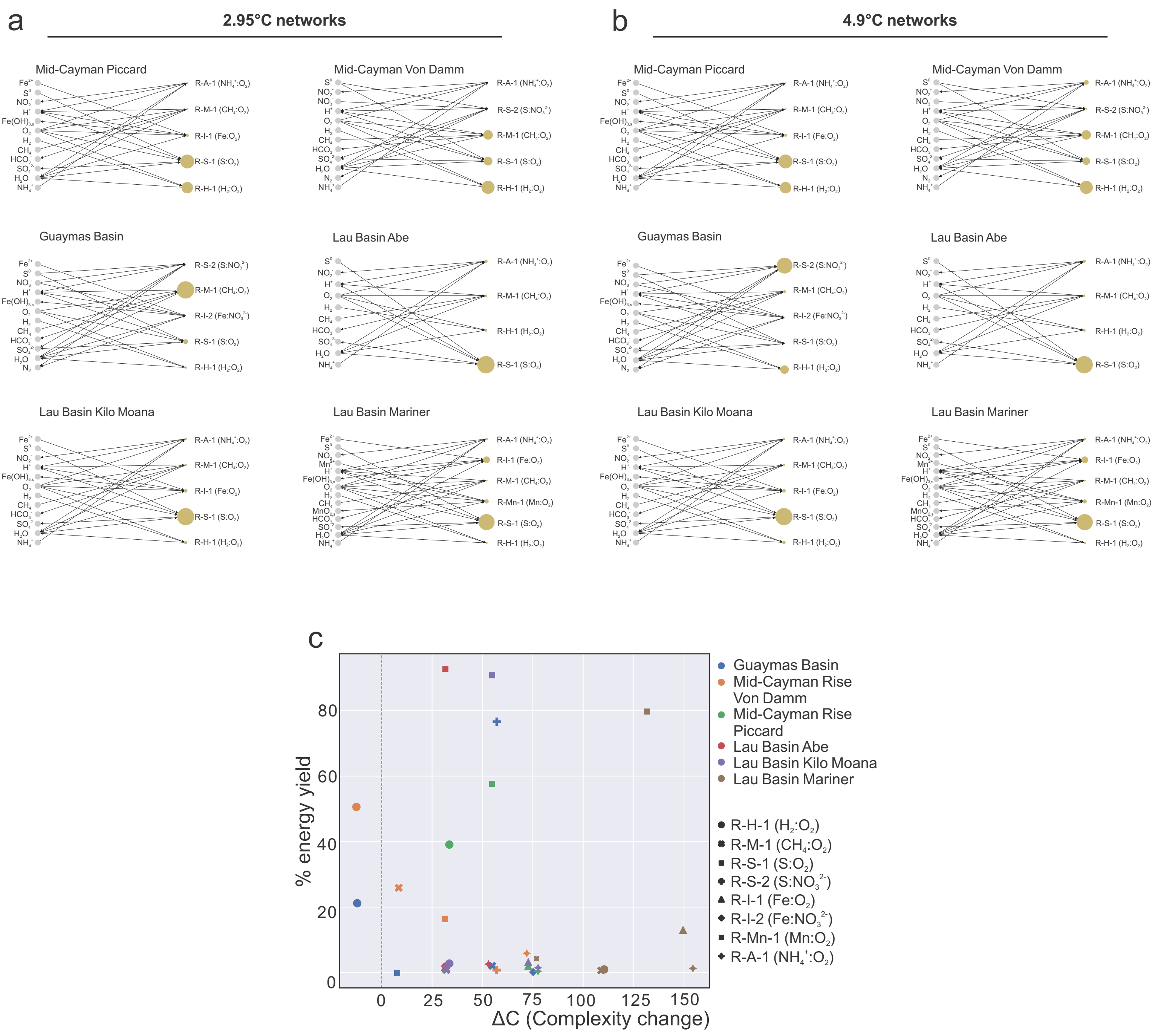


**Supplementary Figure S11. Plume environment networks and network complexity diagram. (a, b)** Plume environment networks based on reactions. The reactions and energy yields for each reaction were based on thermodynamic estimation results at two representative temperatures in plume environments, 2.95°C (**a**) and 4.9°C (**b**). In each network, substrates and products (left side) were connected to each reaction (right side) by an arrow with direction. The size of each reaction was proportional to its energy yield. (**c**) Network complexity diagram representing each reaction's influence on the complexity of the network. In the figure, different colors stand for different hydrothermal environments, different symbol shapes stand for different reactions. The substrates (including electron donor and acceptor) are listed for each reaction in the legend. The x-axis is the change in complexity ($\Delta C$) of the whole network for a node (a rection here) and the y-axis is the percent energy yield of that reaction in the whole community. This network complexity diagram was based on thermodynamic estimation results at 4.9°C.
